# Tension causes structural unfolding of intracellular intermediate filaments

**DOI:** 10.1101/2020.04.22.054379

**Authors:** Frederik Fleissner, Sachin Kumar, Noreen Klein, Daniel Wirth, Ravi Dhiman, Dirk Schneider, Mischa Bonn, Sapun H. Parekh

**Affiliations:** Department of Molecular Spectroscopy, Max Planck Institute for Polymer Research, Mainz, Germany; Institute of Pharmacy and Biochemistry, Johannes Gutenberg-University Mainz, Germany; Department of Biomedical Engineering, University of Texas at Austin, Austin, TX USA

## Abstract

Intermediate filament (IF) proteins are a class of proteins that constitute different filamentous structures in mammalian cells. As such, IF proteins are part of the load-bearing cytoskeleton and support the nuclear envelope. Molecular dynamics simulations have shown that IF proteins undergo secondary structural changes to compensate mechanical loads, which has been confirmed by experimental *in vitro* studies on IF hydrogels. However, the structural response of intracellular IF to mechanical load has yet to be elucidated *in cellulo*. Here, we use *in situ* nonlinear Raman imaging combined with multivariate data analysis to quantify the intracellular secondary structure of the IF cytoskeletal protein vimentin under different states of cellular tension. We find that cells under native cellular tension contain more unfolded vimentin than chemically or physically relaxed specimens. This indicates that unfolding of IF proteins occurs intracellularly when sufficient forces are applied, suggesting that IF structures act as local force sensors in the cell to mark locations under large mechanical tension.

## Introduction

Cell shape and structure result from a well-orchestrated balance of forces acting between the cytoskeleton, extracellular adhesions, and membrane tension^1, 2^. In mammalian cells, the cytoskeleton is the primary load-bearing unit, and it consists of the three filamentous protein networks: actin filaments, microtubules (MT), and intermediate filaments (IFs). While the roles of actin-based networks and MT in cell mechanics and cell function have been extensively studied (reviewed e.g. in^2, 3^), the contribution of IFs to cell mechanics is still relatively opaque, beyond their function in bearing large tensile forces. Of the cytoskeletal proteins, IF constituent proteins have the unique feature of being able to undergo molecular structural changes in response to external loads, at least *in vitro*, as shown by single-molecule experiments and molecular dynamics simulations^4–7^. This structural polymorphism observed outside of a cellular context has led to the hypothesis that IFs could play a role in intracellular mechanotransduction – relaying information about the cell’s mechanical state into biochemical changes that influence cell shape^8–10^ and phenotype^11^.

IF proteins consist mainly of ~45 nm long dimeric building blocks^12^, and crystal structures show that they are at least 67% α-helical^13, 14^. Individual IF proteins self-assemble into networks with fibers consisting of apolar filaments, resulting in the formation of quite diverse networks with location-specific architectures and composition in cells^8^. IFs can form both ionic and hydrophobic molecular interactions resulting in resistant chemical bonds. In addition, proteins such as plectin help bundling and cross-bridging between IFs and to stress fibers and MT ^8, 15^. Vimentin is one IF protein that forms cytoplasmic IF networks and is particularly interesting because of its important role in cell adhesion, mechanical stability^16–18^, and cell migration, as well as intracellular signaling^8, 19^. Specific demonstrations of laser-ablation of super-stretched cells showed that the IF network is load-bearing; however, this effect was not observed on relaxed cells grown on very soft substrates^20^. Moreover, vimentin has been established as a marker of the epithelial-to-mesenchymal transitions (EMT) in embryogenesis and in tumor metastasis, as the cytoplasmic IF network in epithelial cells mostly consists of keratin that is converted to a vimentin-rich IF network during EMT^10, 21–23^.

A complex multi-regime response of the protein structure to deformation has been revealed via computational^7, 24, 25^ and *in vitro* experimental^4, 5^ studies of IF protein mechanics. Researchers have found a hierarchical strain response w by simulating the vimentin central coiled-coil domain under tensile load. The response of the coiled-coil domains to mechanical tension is initially linearly elastic, as only hydrogen bonds are stretched, until the strain exceeds a critical range. For higher strains, hydrogen bonds break, and the coiled-coil α-helical structure sequentially unfolds into anti-parallel β-strand structures^7^. Via this mechanism, each molecule can extend to a multiple of its initial length without breaking. Finally, when no further structural transition is possible to compensate for the load, stretching the covalent peptide bonds and uncoiling of coiled-coil regions causes strain hardening^7, 24^. Besides those structural changes, further extensibility can result from dimeric or tetrameric IF subunits gliding irreversibly against each other^15^. While early work by Astbury and Woods found “dimensional disturbance in the keratin crystallite”^26^, the more recent studies by Fudge et al.^27^ on hagfish slime and Pinto et al.^28^ on human vimentin gels used X-ray diffraction and wide-angle X-ray spectroscopy to show that β-sheet signatures appeared in tensed samples. Infrared spectroscopy on horse hair, consisting of mostly keratin IF proteins, which can be crosslinked by disulfide bonds into “hard”keratins^8^ also showed an increased contribution from β-sheets under tensile strain^29^. While these studies clearly showed structural transitions of hard keratins and vimentin under tension *in vitro*, it is unclear if such transitions occur within the intracellular vimentin IF network, where the cellular environment has been shown to modify cytoskeletal properties and mechanics^30^.

Few experimental methods allow analyzing protein structure within cells (reviewed in ^31^ and ^32^). Förster Resonance Energy Transfer (FRET) sensors can be used to selectively measure intramolecular distances that can be correlated to tension acting on a target protein^33, 34^. However, this approach is challenging to realize in bundle-forming filaments where cross-talk between neighboring dye molecules complicates the interpretation of the FRET signal^35, 36^. Cys-shotgun labeling of exposed cysteines is another method, introduced by Discher and colleagues, to show that IF proteins undergo conformational changes in response to cell tension by exposure of otherwise buried cysteines^37, 38^. However, both FRET measurements and Cys-shotgun labeling are unable to probe a protein’s secondary structure, which is vital to rationalizing if the *in vitro* load-bearing mechanism involving structural transitions is relevant for intracellular IF networks.

Protein secondary structure can be probed in a noninvasive way using vibrational spectroscopies, such as infrared or Raman spectroscopy^39^. Depending on the local hydrogen-bonding pattern (i.e. secondary structure), the atoms in the protein backbone (C=O vibrations in peptide bonds) show molecular vibrations at different vibrational frequencies, resulting in shifted Raman peaks in the Amide I region of the vibrational spectrum^40–42^. This link between protein structure and spectral response can be exploited to determine the average secondary structural composition within the probed volume with high accuracy using quantitative spectral decomposition^40^.

In this study, we investigate if tension-induced secondary structural transitions of vimentin IF can be observed in adherent cells using quantitative, hyperspectral broadband coherent anti-Stokes Raman scattering (BCARS) vibrational microscopy with sub-micrometer spatial resolution. BCARS microscopy combined with isotope-substitution and a multivariate spectral analysis protocol is used to resolve the vimentin secondary structure within cells. By exploiting the unique spectral features from isotopically-labeled vimentin to distinguish the vimentin Raman signal from the cellular protein background, we isolate the spectral fingerprint of deuterated vimentin in cells and determine its secondary structure for cells in different tensile states. Our results show that cellular tension is sufficient to alter the secondary structure of intracellular vimentin IFs, resulting in more β-sheet content with increasing tension.

## Results

Since mechanical deformation has previously been shown to cause structural transitions in IF-composed structures (hagfish slime, vimentin gels, and hair) *in vitro*^27–29^, we first tested how mechanical deformation affected the secondary structure of vimentin IFs when it was polymerized *in vitro*. Using a micromanipulator on our nonlinear Raman (BCARS) microscope (see methods for details), we acquired BCARS spectra from relaxed and physically tensed vimentin *in vitro* (**Figure 1A and B**). To obtain a sufficiently strong Raman signal from a single position it was necessary to measure vimentin bundles. Upon visual inspection of the Amide I region of the Raman-like (RL) spectra (**Figure 1C**), which is sensitive to the protein secondary structure, clear spectral changes were observed between the relaxed and tensed vimentin IF bundles (**Figure 1C**, see also SI **Figure S1** for additional measurements). Spectral decomposition of the Amide I region using known peak shapes and locations for α-helices, β-sheets, and random coil (rc) regions allowed for calculation of the average secondary structure^40, 43^. The relaxed vimentin IF bundles – with a sharp peak at ~ 1650 cm^−1^ – showed a predominantly α-helical structure (64% α-helix, 15% β-sheet and 21% rc) after decomposition, which compares favorably to circular dichroism data from Meier *et al.*^44^. On the other hand, tensed samples – with a broadened (more disordered) and blue-shifted Amide I band, revealed an increased β-sheet structure and reduced helicity (40% α-helix, 32% β-sheet, 27% rc), consistent with the observation of a banding pattern originating from β-sheet observed by Kreplak and coworkers^45^.

**Figure 1:**
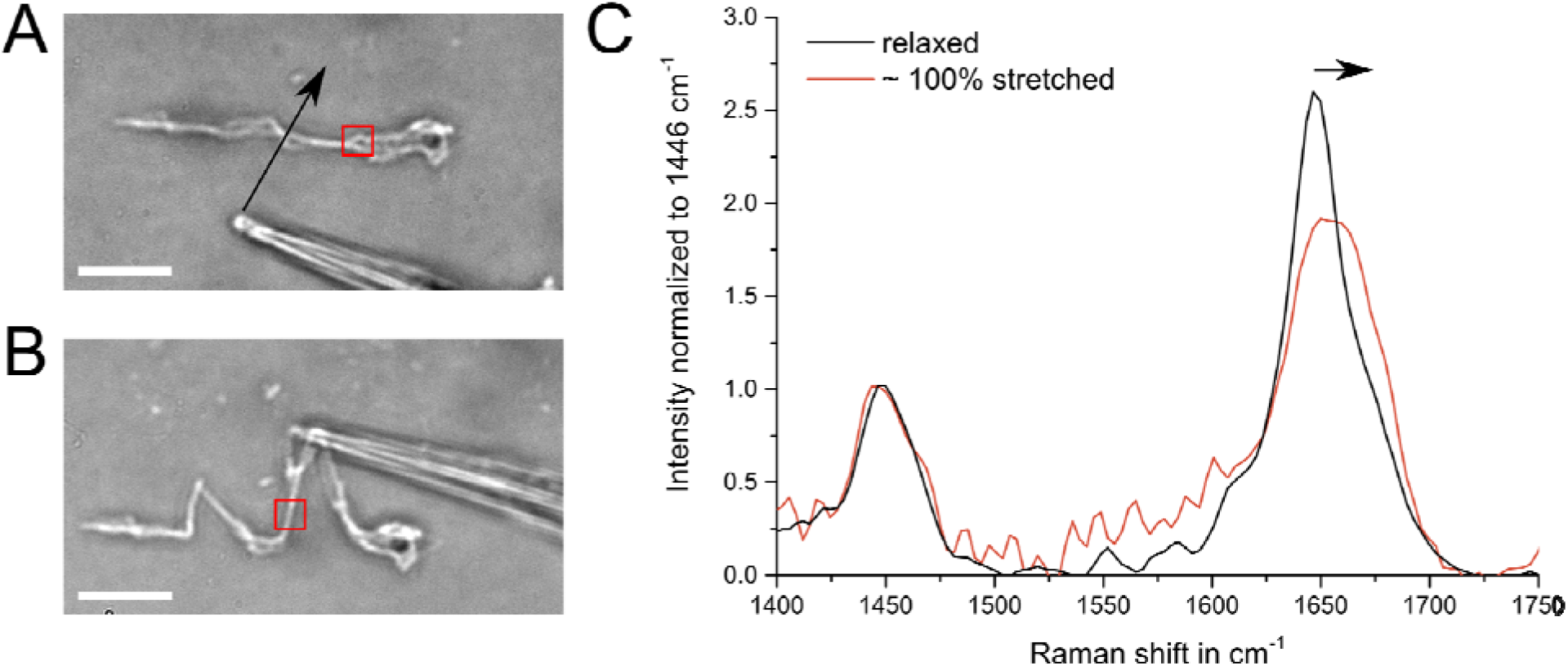
*In vitro* deformation experiments of Vimentin hydrogels show protein unfolding. **A)** Bright field microscopy image of a polymerized vimentin IF filament bundle adhered to a coverslip. A glass capillary mounted on a micromanipulator was threaded below the filament and moved along the plane as indicated by the arrow for pulling. **B)** The filament bundle stretched out under tension, as was apparent from the bending of the capillary tip. Scale bars in both images represent 10 μm, and the filament bundle diameter was found to be ~0.5 μm. The filament regions probed by BCARS are indicated by red squares. **C)** RL spectra of relaxed and more than 100% strained vimentin bundles showing clear changes in the Amide I region (1600-1680 cm^−1^) from a narrow band at 1640 cm^−1^ (assigned to α-helical structure) to a blue-shifted and broadened band indicating the presence of additional β-sheet and random coil structure under load. Spectra were normalized to the CH_2_ deformation mode at 1446 cm^−1^. For the relaxed vimentin, an average peak center of 1645 ±1.6 cm^−1^ was found while for the pulled vimentin samples the center was 1651 ±1.0 cm^−1^. A two-sample t-test assuming unequal variance resulted in a significant difference. Further experimental data can be found in the supplementary information (**Figure S1**).

Changes in the Amide I band in purified, reconstituted vimentin IF networks under tension could be unequivocally attributed to changes in vimentin structure under strain; however, attributing a particular Amide I spectral change as coming from a specific protein within a cell is extremely challenging. Cells contain thousands of different proteins with a total protein concentration of ~200 mg/mL^46^, and each protein contains very similar (and overlapping) vibrational moieties. Thus, the BCARS spectra from any location inside a cell will consist of the sum of all proteins within our excitation volume (0.4 x 0.4 x 5 μm^3^), making it nearly impossible to produce vibrational fingerprints from a *specific* protein in the cell. It is therefore necessary to separate the contribution of the target protein, in this case vimentin, from that of the remaining cellular background. A convenient way to create spectral contrast with minimal perturbation is stable isotope substitution, such as when hydrogen (^1^H) is replaced by deuterium (^2^H, or D), as is commonly done in NMR spectroscopy.

We produced recombinant, isotopically-substituted vimentin using *E. coli* grown in M9 medium containing deuterated carbohydrates as the only carbon source (see Methods). This resulted in partial deuterium-hydrogen exchange for all amino acids in the protein, as confirmed by mass spectrometry (see SI, **Figure S2**). In the case of 100 % D-H exchange, the molecular weight of the deuterated vimentin would be at most 3.7 kDa more than the native human vimentin protein, including the His-tag, which had a molecular weight of 55 kDa). Purified D-Vim had a similar band to H-Vim in SDS-PAGE (see **Figure 2A** and SI **Figure S3**), and perhaps more importantly, the functionality of the protein was confirmed by successful *in vitro* polymerization.

**Figure 2:**
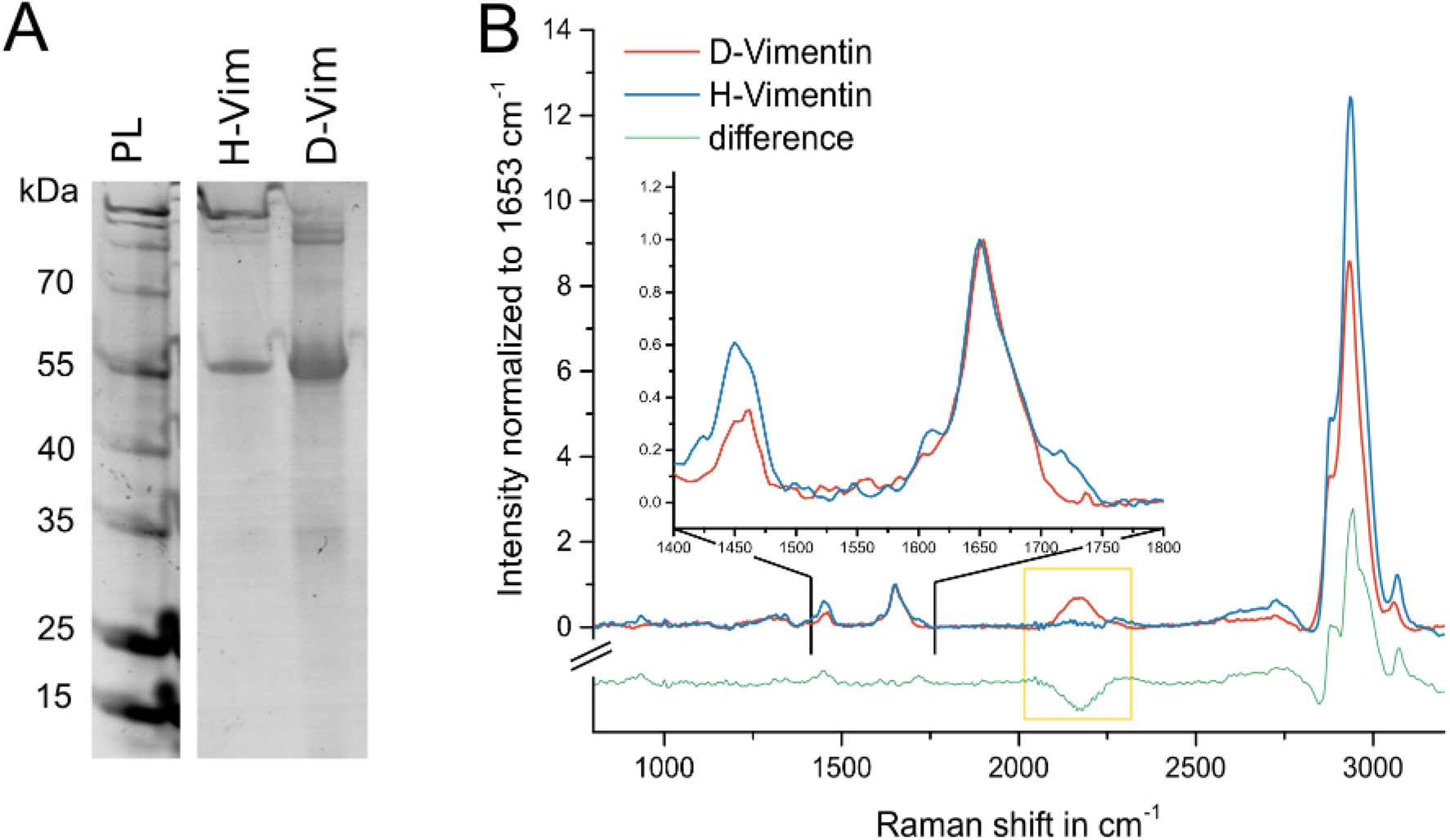
Characterization of deuterated Vimentin. **A)** Coomassie-stained SDS-PAGE gel of native vimentin (H-Vim) and vimentin produced from deuterated acetate (D-Vim) showing similar molecular weight at 55 kDa. The first lane shows the protein ladder with molecular masses of the ladder given on the left. **B)** RL spectra of native vimentin and D-Vim. Spectra were normalized to the peak intensity at 1653 cm^−1^. For D-Vim the RL spectra show an additional vibrational band centered at 2150 cm^−1^ originating from the CD modes. At the same time, the CH band (2800-3100 cm^−1^) and the CH_2_ deformation mode at 1450 cm^−1^ had a reduced intensity compared to native vimentin as indicated in the difference spectrum. The Amide I band, reflecting the protein C=O backbone vibration, was nearly identical for both proteins.

**Figure 2B** shows the RL spectra of normal vimentin (H-Vim) and deuterated vimentin (D-Vim) in polymerized vimentin fibers. The additional peak at 2150 cm^−1^ in D-Vim showed the unique carbon-deuterium (CD) stretch spectral feature in the so-called vibrational quiescent region, which – as we show below – allowed for uniquely identifying signals coming from D-Vim compared to the intracellular cytosolic protein pool. Moreover, the Amide I region (1550-1700 cm^−1^) appeared almost identical for D-Vim and Vim, indicating: 1) that hydrogen-deuterium exchange on NH groups in peptide bonds was minimal or the ND rapidly exchanged in our hydrogen-based buffers and 2) the secondary structure of D-Vim was the same as H-Vim. We note that even though D-Vim was produced recombinantly from a nearly 100 % deuterated carbon source, deuterium incorporation into the protein was clearly not 100 %. Strong CH signals from the protein were still observed in the RL spectrum of D-Vim (**Figure 2B, red**). The observed carbon-hydrogen bonds in D-Vim likely result from hydrogen-deuterium exchange during protein production and purification since we used H_2_O-based M9 growth medium and buffers.

To investigate spectral features of vimentin intermediate filaments in cells, we incorporated recombinantly produced D-Vim into HeLa cells following the protocol illustrated in **Figure 3A-D**. HeLa cells were chosen as a model mesenchymal cell as they are vimentin expressing, uncomplicated in handling, and well-suited for the lentiviral transfection used below^47^. First, the native IF network was depolymerized by addition of cycloheximide (see methods and SI **Figure S4** and **S5**)^48^. Then, recombinantly produced D-Vim monomers were microinjected into the cells (D-Vim = 1.8 mg/mL) and re-formation of the IF network was induced by replacing the media with normal (cycloheximide-free) media^49^. After 12 hours of incubation in normal growth media, the IF network recovered without any obvious accumulation of D-Vim puncta in the cytosol, and D-Vim was successfully co-polymerized into the intracellular vimentin IF network (see SI **Figure S6**).

**Figure 3:**
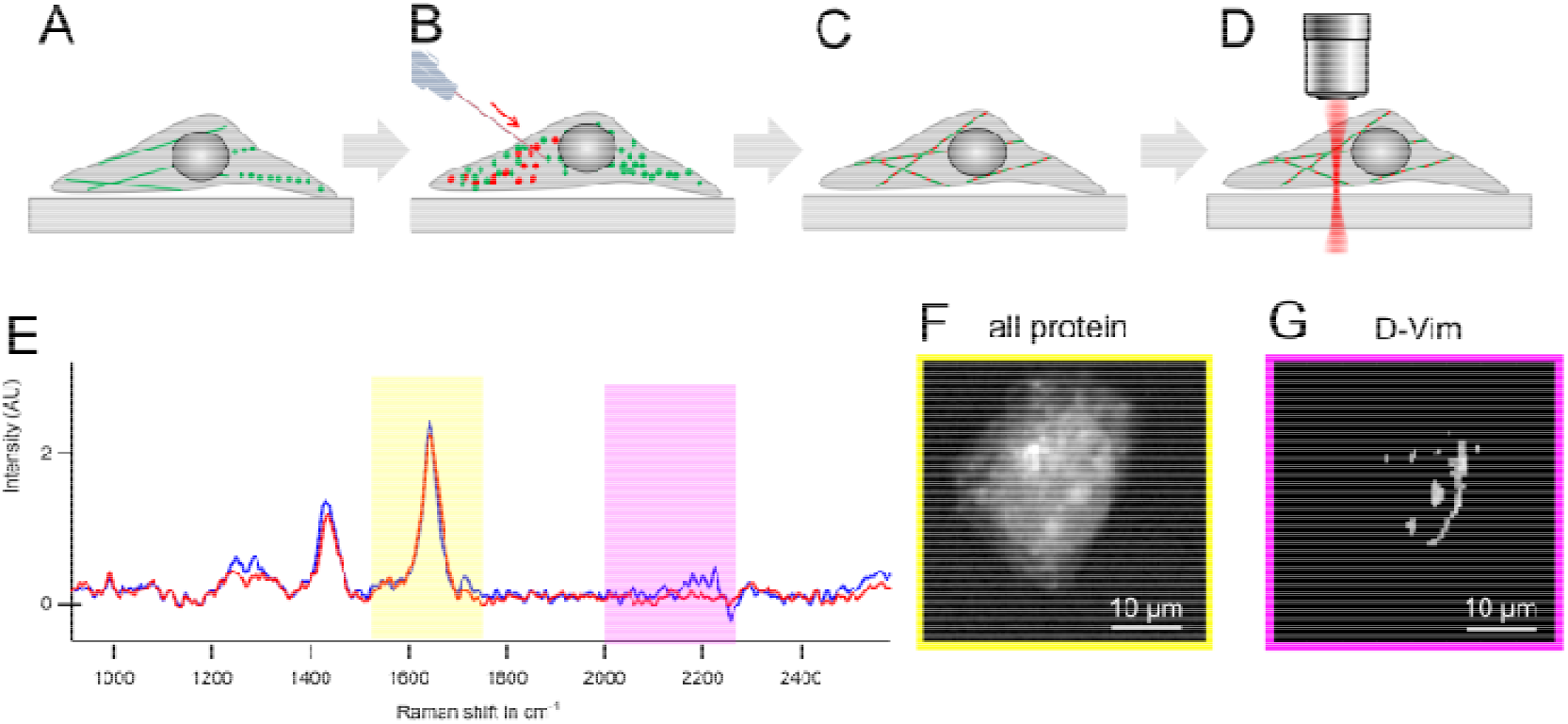
Deuterated vimentin creates label-free contrast within cells. The native IF network of adherent HeLa cells (depicted in green) was depolymerized by treatment with cycloheximide **(A)** and deuterated vimentin concentration (1.8 mg/ml, red) was injected **(B)**. After incubating overnight, D-Vim incorporated in the reformed IF network **(C)**, and cells were fixed for hyperspectral BCARS microscopy to obtain local chemical information **(D)**. **E)** RL spectra obtained from two different cell locations show differences in the silent (purple) region resulting from additional molecular vibrations from the CD bonds of D-Vim. **F)** The total protein distribution within a cell measured by the integrated intensity of the Amide I band (yellow region in E) in each spatial pixel. **G)** The D-Vim distribution for the same cell as in F is shown as the integrated intensity over CD region (purple region in E).

Following this protocol resulted in a cellular vimentin network containing D-Vim as well as H-Vim, and cells should show a CD vibration in the silent region of the RL spectra. This vibration could, in principle, be used to localize D-Vim in the cell. A similar approach using CD isotope substitution has already been used to identify newly synthesized protein in cells^50, 51^ and for tracking cell-penetrating peptides in cells^5, 53^. We observed the CD vibration (at ~2200 cm^−1^) from D-Vim when it was sufficiently concentrated in cellular IF fibers (**Figure 3E**). This signal could be used to create contrast showing the concentration of D-Vim throughout the cell, which is clearly different from the distribution of all proteins in the cell (**Figure 3F and G**).

We combined the ability to incorporate D-Vim into the cellular IF network with our previously developed analysis protocol to isolate the spectral response of isotope-labeled proteins in cells^53^ in order to determine how mechanical perturbations affected the intracellular vimentin secondary structure. The cellular mechanical state was manipulated both physically and biochemically by culturing cells on soft substrates and by drug treatments that interfere with cell-generated tension, respectively. Mesenchymal cells, such as HeLas, grown on (infinitely rigid) glass substrates are known to have a cytoskeleton that is under increased tension or pre-stress compared to when grown on soft substrates ^54, 55^. Confocal microscopy of HeLa cells expressing GFP-tagged vimentin showed that the IF network was laterally sprawling and occupied a large area, and cells showed a typical fried egg-like appearance when grown on collagen-coated glass (**Figure 4A and B**). Vimentin filaments formed a perinuclear cage and spread out toward the cell periphery.

**Figure 4:**
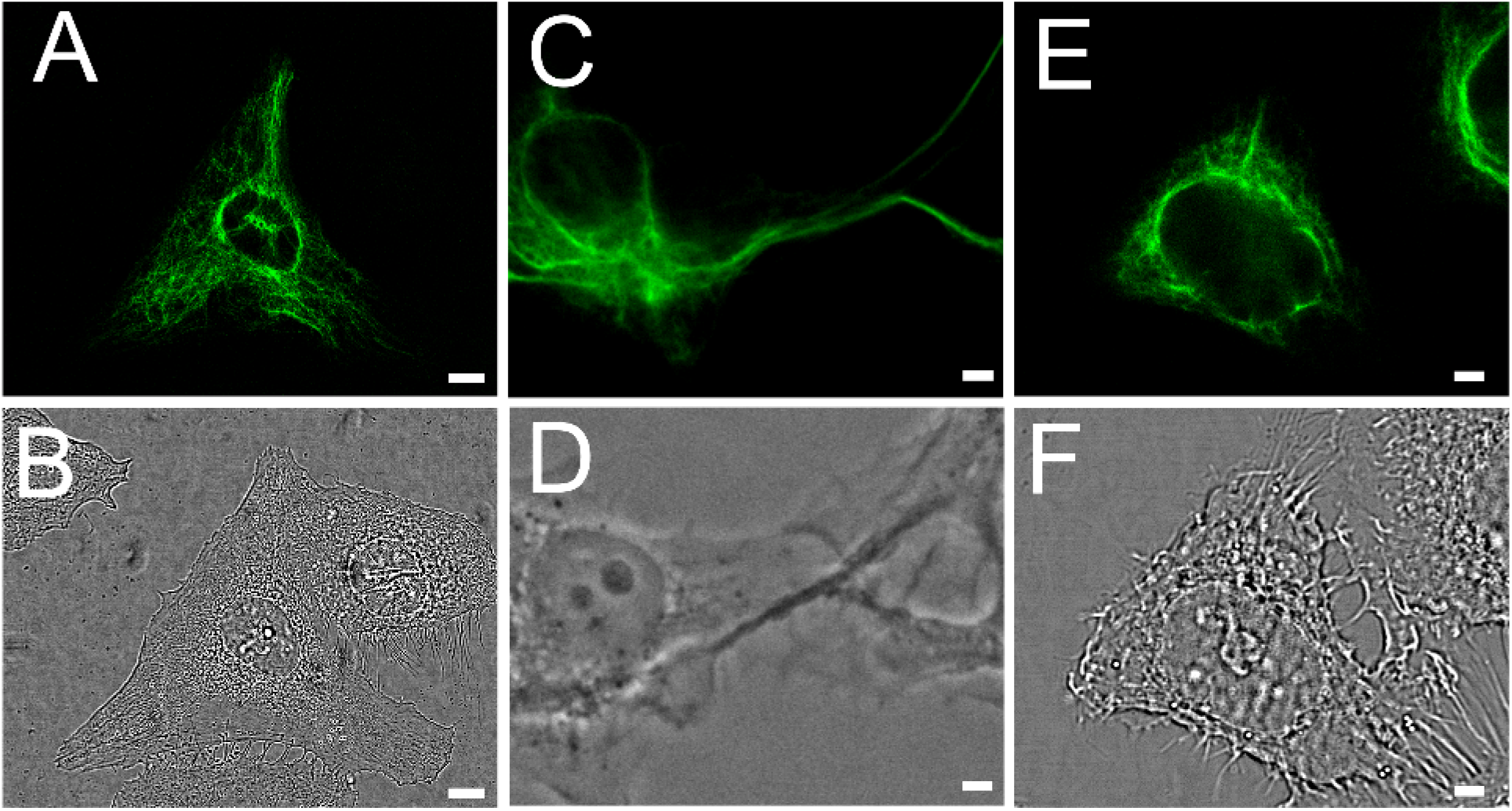
The architecture of the IF network is altered by substrate stiffness and drug treatment. Confocal microscopy images of HeLa cells expressing GFP-tagged vimentin (**A, C, E**), and brightfield confocal images showing cell morphology (**B, D, F**). **A, B)** (Control) Cells grown on collagen-coated glass substrates were spread out with a broad vimentin network. **C, D)** Blebbistatin treatment of cells grown on collagen-coated glass caused cells to round up and shrink, and the vimentin IF network became less sprawling and more bundled. **E, F)** Cells grown on a collagen-coated soft substrate (0.1 kPa polyacrylamide) rounded up compared to control cells (A, B) and showed a collapsed vimentin IFs only as a perinuclear bundled network. All images were taken from imaging planes below the nucleus. Scale bars represent 10 μm.

Intracellular tension is produced by myosin-based contraction of actin stress fibers (in two-dimensional cell culture), and to investigate the effect of cell-generated tension on vimentin IFs, we treated cells with blebbistatin, a non-muscle myosin II inhibitor. The drug inhibits ATP hydrolysis and thereby blocks acto-myosin contraction^56^, resulting in relaxation of cell-generated tension. **Figures 4C and D** show HeLa cells grown on glass substrates treated with blebbistatin. Treated cells partially rounded up, a hallmark of non-muscle myosin II contraction inhibition, compared to those cultured without blebbistatin. The vimentin IFs (**Figure 4C**) appeared wavy and less sprawling than before the treatment but still spanned large distances within the cytosol.

From previous studies, it is also known that substrate stiffness regulates cellular tension^57, 58^. As an alternative to chemically reducing the tension, we cultured cells on collagen-coated soft substrates (0.1 kPa polyacrylamide gels) to again reduce the cell-generated tensile forces. HeLa cells exhibited an altered morphology when cultured on soft substrates, with a more perinuclear vimentin network and compact shape (**Figure 4E and F**) compared to the cells grown on glass (**Figure 4A and B**).

The influence of cellular tension on the vimentin secondary structure was analyzed by comparing the shape of the Amide I vibration in D-Vim RL spectra from cells that were grown on collagen-coated glass (control) to cells in which tension was physically or chemically relaxed. For each group, we acquired hyperspectral BCARS images of individual cells. Each data set covered an area of 40 x 40 μm^2^, containing at least one cell, with a step size of 0.5 μm per spatial pixel. After retrieval of the RL spectra, the data were thresholded to retain pixels showing sufficient CD intensity. Spectral components for D-Vim, as we l as the cell background, were derived via the multivariate curve resolution (MCR) approach, using the CD vibration as a handle, as previously established^53^ (see methods for a detailed explanation). Exemplary spectra from this process for the D-Vim and cell background component spectra can be found in the supplementary information (see SI **Figure S7-S11**).

The spectra in **Figure 5A** show the Amide I region of spectrum averaged over an entire cell grown on collagen-coated glass, the average MCR-derived, D-Vim component from all cells that were grown on glass, and the spectrum from relaxed, *in vitro* polymerized vimentin fibers. **Figure 5B** shows single-ce l spectra for the MCR-derived, D-Vim component, the MCR-derived cell background component, and the spectrum calculated by averaging all spectra over a single cell grown on collagen-coated glass. The CD band (~ 2200 −2400 cm^−1^) was considerably larger in the MCR-reconstructed spectrum for D-Vim (**Figure 5B, red**) compared to the cell background or cell-averaged spectrum, confirming the specificity of the component for deuterated vimentin. The Amide I region of the D-Vim component spectrum extracted from cells grown on glass (**Figure 5A, red**) showed a clearly different spectral fingerprint compared to *in vitro*, polymerized vimentin (**Figure 5A, yellow**). While the CH_2_ stretching vibration at 1450 cm^−1^ had the same peak location and width in both cases (as one would expect), the shape and maxima of the Amide I band were both altered in cells grown on glass. Changes in the Amide I band shape of the MCR-derived, D-Vim component show different structural compositions of D-Vim in cells compared to polymerized *in vitro*. The peak maximum for the Amide I band was shifted from 1650 cm^−1^ for *in vitro* polymerized vimentin to 1663 cm^−1^ for the intracellular D-Vim MCR-derived component spectrum, indicating that intracellular vimentin contained more β-sheets while the *in vitro* polymerized vimentin was more helical. The peak maximum of the Amide I region in the average cell spectrum (**Figure 5A, blue**) coincided with that of the D-Vim component.

**Figure 5:**
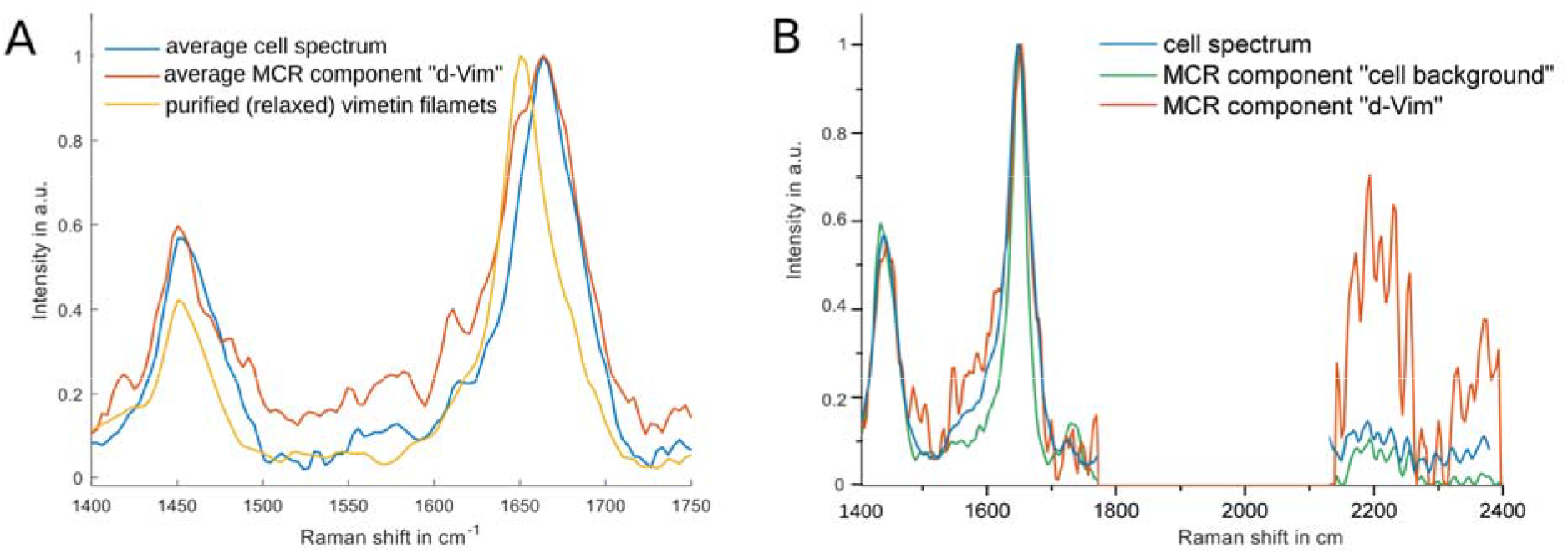
Separation of the vimentin component by multivariate curve resolution. **A)** Spectra showing the Amide I region of the entire cell (blue) and the MCR-derived, D-Vim component averaged over the group of cells grown on collagen-coated glass without further treatment (red). For comparison, the average spectrum from pure vimentin polymerized *in vitro* (yellow) is depicted. **B)** A representative average cell spectrum (blue) and derived MCR components assigned to the cell background (green) and D-Vim (red) are shown measured with a single cell grown on collagen-coated glass.

Average spectra of each MCR-derived component (cell background and D-Vim) for each cell group (control (glass), soft gel, and blebbistatin-treated) are shown in the SI (see SI **Figure S10-S13**). All D-Vim components clearly showed a dominant CD vibration in the D-Vim component (and the CD vibration was substantially smaller in cell background component). To determine if cellular tension affected the structure of vimentin within cells, we compared the average MCR-derived, D-Vim Amide I spectrum from cells treated with blebbistatin or cell grown on soft substrates to control cells grown on collagen-coated glass (see SI **Figure S14 and S15**). Treatment with blebbistatin resulted in a red-shifted, narrower Amide I band that was centered at 1652 cm^−1^(**Figure 6A**), indicative of a more α-helical (native) structure and lower contributions from β-sheet structures compared to control cells grown on collagen-coated glass. In addition, the D-Vim component spectrum from cells treated with blebbistatin had a narrower Amide I region compared to the D-Vim peak from cells on glass (**Figure 6A**), similar to the spectrum of purified vimentin *in vitro* (**Figure 5A** (yellow)).

**Figure 6:**
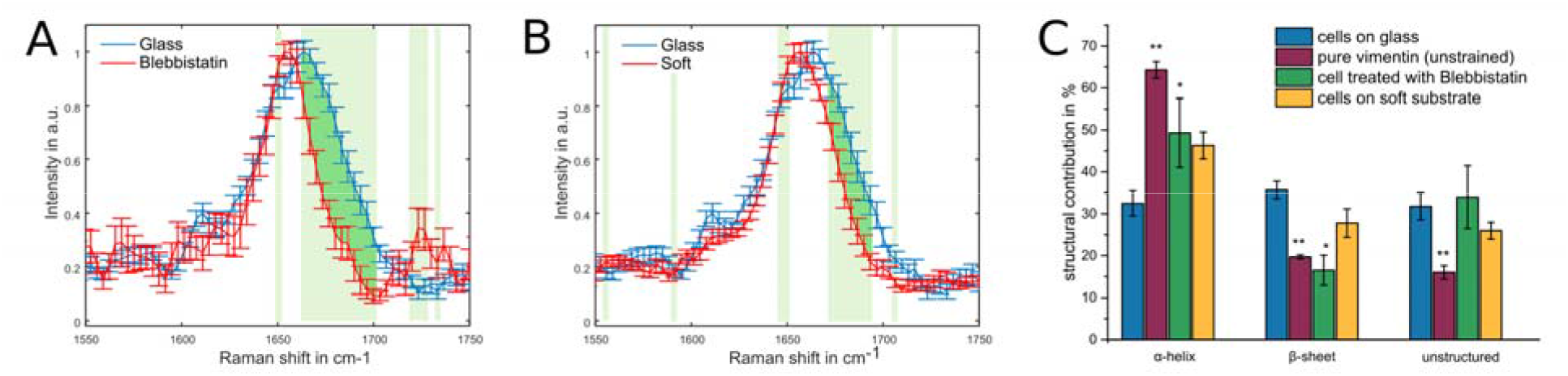
The vimentin structure is affected by cellular tension. Spectral analysis of cell grouped by different intracellular stresses showed changes in the Amide I region demonstrating differences in vimentin structure. **A)** Average of the Amide I bands from MCR-derived, D-Vim spectra of cells grown on collagen-coated glass substrates (blue, n=14), and after blebbistatin treatment (red, n=7). **B)** Average of the Amide I bands from MCR-derived, D-Vim spectra of cells grown on collagen-coated glass substrates (blue, n= 14), and cells grown on collagen-coated soft substrates (red, n=17). In both panels A and B, error bars are standard error of the mean (SEM) for each data point and statistically significant different spectral regions are highlighted by light green boxes. Dark green shades highlight the reduction in β-sheet and random coil structure. **C)** Secondary structure content of vimentin derived via spectral decomposition of the Amide I band (either from pure vimentin filaments or the MCR-derived component). Error bars are standard deviation from the stated number of samples. Statistically significant differences between cells cultured on collagen-coated glass and other groups from two sample t-tests are indicated as (*) for P ≤ 0.05 and (**) for P ≤ 0.01.

Cells that were cultured on soft substrates showed a weaker intensity at the right shoulder of the Amide I band and significantly lower peak intensities around 1680 cm^−1^ with respect to cells from glass in the D-vim component spectra (**Figure 6B**). The Amide I center frequency of 1654 cm^−1^ was similar to that of blebbistatin-treated cells. These changes relative to the D-Vim spectra from cells cultured on glass again indicate that vimentin is more natively (α-helically) structured when cultured on soft substrates. Comparing the MCR-derived, D-Vim component spectra for cells from each sample group, we find statistically significant differences (P < 0.05) when comparing cells with reduced tension (either via treatment with blebbistatin or via culturing on soft substrates) against those grown on glass with “native” cytoskeletal tension (**Figure 6A and B** light green boxes).

In addition to the physical and chemical tension reducing methods already shown, we employed a blunter method of cellular tension reduction via treatment with latrunculin-A, a marine toxin that promotes depolymerization of the actin cytoskeleton^59, 60^, for cells cultured on collagen-coated glass. The Amide I spectra for latrunculin-A treatment (see **Figure S16**) resemble the results for soft gels, again showing that tension reduction results in more helical (native) vimentin in cells.

In order to quantify the altered Amide I line shape and relate these changes to the D-Vim protein structure, we decomposed the MCR-derived, D-Vim Amide I band from each cell using our previously established Amide I spectral decomposition^53^. **Figure 6C** shows the resulting average contributions (and standard deviations) of α-helical, β-sheet, and random coil structural component for all cell groups, as well as for *in vitro* polymerized vimentin. The spectral decomposition can be found in the SI **Figures S13** and **S15**. While *in vitro* polymerized vimentin was found to be predominantly α-helical (64%) when measured by BCARS, consistent with previous results^44^ and our own circular dichroism measurements (see SI **Figure S17**), the amount of helical structure was only 33% for intracellular vimentin in control cells. With a lowering of the α-helical content in cells, the calculated β-sheet content showed the opposite trend – being 36% for cells on glass compared to 20% for *in vitro* vimentin. Both chemical and physical relaxation of cells resulted in an intracellular structure of vimentin that was significantly more α-helical (and contained fewer β-sheets), similar to *in vitro* vimentin. We found that for cells treated with blebbistatin, vimentin was on average 49% α-helical and 17% β-sheet structured while for cells grown soft substrates the structural composition resulted in 46% α-helix and 27% β-sheet content. The amount of unstructured vimentin was found to be statistically similar for all cell groups, indicating that the relaxation of cell tension did not increase random coil amounts in intracellular vimentin IFs. However, the amount of random coil for cytosolic vimentin was significantly higher for all cell groups than for *in vitro* polymerized vimentin.

Previous work has shown that the level of phosphorylated vimentin, particularly at the Serine 55/56 residue, correlated with cell tension in such a way that phosphorylated vimentin in IFs was reduced with a higher tensile state^61, 62^. As the intracellular vimentin secondary structure was shown to be different for HeLa cells grown on different stiffness substrates, we hypothesized that this might also result in phosphorylation changes in vimentin IFs. Therefore, vimentin organization and level of vimentin phosphorylation at serine 55 (pSer55) in response to the mechanical properties of substrate were studied using immunofluorescence as shown in **Figure 7**. HeLa cells expressing GFP-vimentin show a vimentin network structure on both soft (3 mg/ml) collagen hydrogels and stiff, collagen-coated glass surfaces (**Figure 7A and 7D**). For the soft hydrogel surface, we observed a fragmented vimentin network and pellet-like vimentin structures (indicated by blue arrows in **Figure 7A**). On the other hand, a highly assembled and mature vimentin filament network was observed on stiff, collagen-coated glass surfaces (**Figure 7D**). **Figures 7B and 7E** show immuno-fluorescence images of pSer55 for the same cells shown in **Figure 7A** and **7C**, respectively. The merged channels of GFP-Vim and Vim-pSer55 depicted in **Figure 7C** and **E** resemble each other in their spatial distribution and confirm the selective staining of vimentin only when the cells are cultured on soft substrates, where the tension is low and the vimentin network is predominantly α-helical (**Figure 7G**). Images of cells on additional collagen gels (0.1 mg/ml and 1 mg/ml) are shown in the SI (**Figure S18**), confirming that increased pSer55 is found in vimentin in cells cultured on soft substrates.

**Figure 7:**
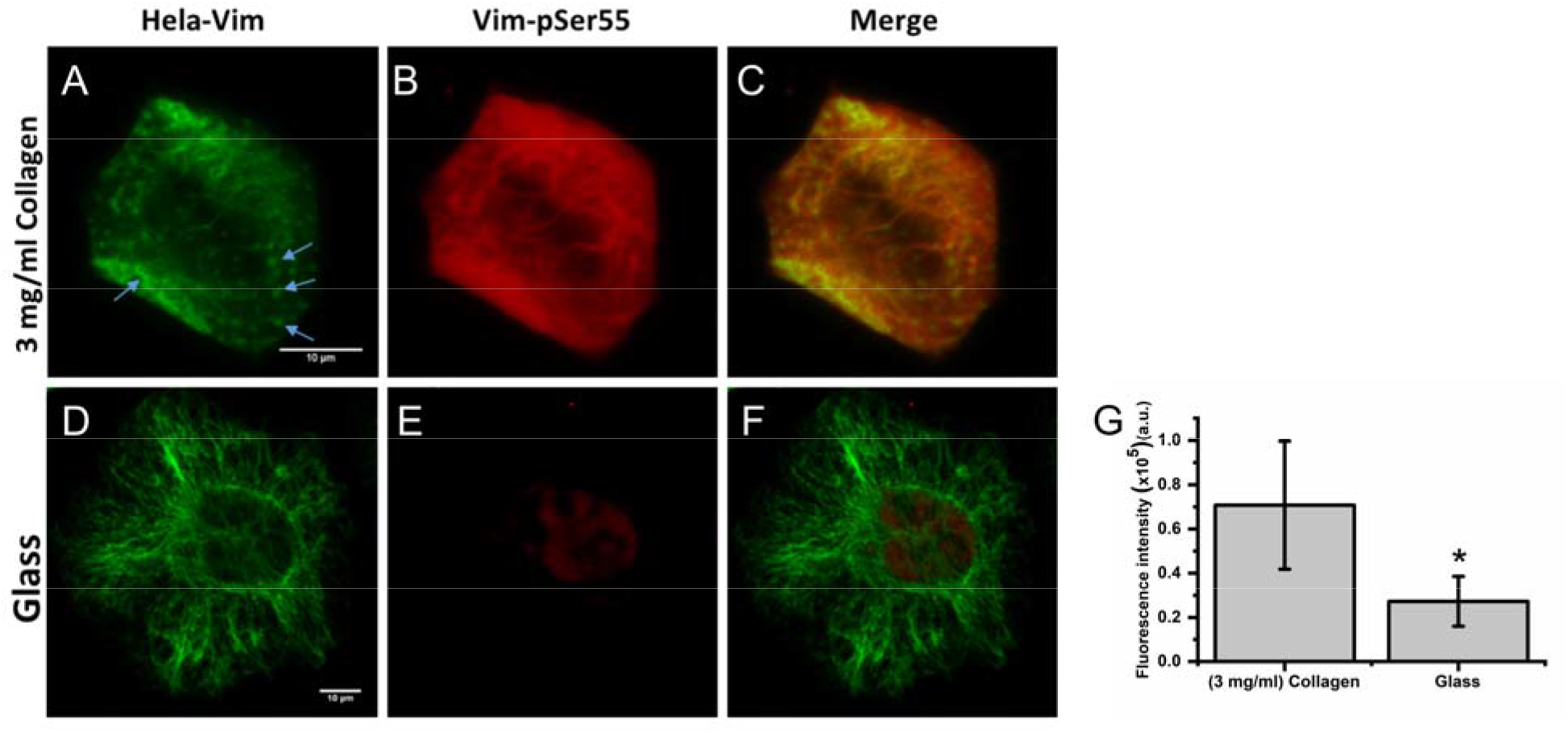
Vimentin phosphorylation is decreased by cellular tension. Immunofluorescence imaging of cells grown on soft collagen hydrogels and collagen-coated glass surfaces shows GFP-tagged vimentin **(A, D)**, pSer55-antibody **(B, E)** and merged **(C, F)**. Scale bars represent 10 μm. Quantification of pSer55 fluorescence intensity showed a significant increase in fluorescence intensity for cells grown on soft collagen substrates **(G)**, indicating high-phosphorylated vimentin.

## Discussion

In this work, we presented *in situ* measurements of the vimentin secondary structure in human cells under different cellular tensions. Pulling on purified and *in vitro* polymerized vimentin filaments adhered to a coverslip resulted in a clear change in its Raman signature, showing a conformational change under load. This is consistent with previous results from MD simulations^7, 24, 25^ and force-strain experiments^4, 5^. As unfolding has been purported to start at ~20% strain^24^, the applied strain of more than 100% of the initial filament length likely caused most vimentin molecules in the stretched vimentin bundle to undergo a transition from an α-helical to a β-sheet (and even into a random coil) structure.

The influence of cell-generated mechanical tension on the vimentin secondary structure was measured after modulating the mechanical state of cells both chemically and physically and using BCARS vibrational imaging of microinjected, deuterated vimentin (D-Vim) that incorporated into an intracellular vimentin network. Employing our previously developed chemometric spectral analysis protocol^53^ yielded not only a unique D-Vim component, but also a “cellular background” component of non-D-Vim proteins from the same spatial pixels in each cell. We found that the average cellular spectrum (see SI **Figure S9**) and the cellular background (non-D-Vim) component for cells under all conditions (see SI **Figure S13A**) were largely statistically identical – most importantly in the Amide I region (see SI **Figure S13B and C**), indicating that the D-Vim component we extracted accurately reflects the D-Vim Amide I spectral fingerprint. Noteworthy, as the data processing route used here is based on unsupervised multivariate separation without further training data, it is possible that spatially correlated molecules (to D-Vim) could also contribute to the extracted D-Vim spectral component. This is highly unlikely, as it would need to be the same molecules in the cases of cells cultured on soft substrates, cultured with blebbistatin, and cultured with latrunculin-A, since these all showed very similar changes in the D-Vim component.

From our spectral decomposition of the extracted D-Vim component we found that control cells grown on collagen-coated (rigid) glass substrates, i.e. cells that are highly tensed, exhibited a D-Vim component that was structurally distinct compared to (relaxed) *in vitro* polymerized vimentin (**Figure 5A**). Furthermore, our data show that a reduction of cellular tension by either reducing the substrate stiffness or deactivating traction force generation through blebbistatin (or latrunculin-A) inhibition of acto-myosin contraction led to a more native (α-helical) D-Vim component spectrum (**Figure 6A and B**). It was previously demonstrated that *in vitro* IF filaments undergo an α-to-β transition under tensile load^4, 27^. Our result, that the secondary structure of native vimentin IF filaments in cells is partially unfolded by cell-generated tensile force, is coincident with substantial reduction of vimentin phosphorylation. This indicates that a vimentin force-response mechanism also occurs for cells grown on coverslips, not only under extreme conditions, e.g. large external deformation.

Our findings are in line with previous studies that used blebbistatin as a selective drug only inhibiting myosin but not affecting IF or microtubules^63, 64^ but contrast with blebbistatin-induced depolymerization of vimentin IFs as reported by Johnson *et al.*^37^. We did not observe blebbistatin-induced depolymerization of vimentin in cells or *in vitro* (see SI **Figure S19**). Importantly, our result showing more helical (natively folded) vimentin structure in blebbistatin-treated cells was further underscored by a similar finding when culturing cells on soft substrates and cells treated with latrunculin-A, which are both well known to reduce traction forces in adherent cells^59, 65^. The consistency of these experiments points to the idea that cell-generated tension puts intracellular vimentin IFs under large forces, resulting in an altered secondary structure of the protein in the IF network.

The unfolding of coiled-coil regions in proteins has been proposed to be a fundamental force-response mechanism in nature^6, 66^. Such a mechanism endows a material with the tunable elasticity of being soft at low tensions while becoming nonlinearly stiffer under larger tensile forces (at large deformations).

Previous work on fibrin, the blood clot forming protein, revealed features similar to those observed for vimentin: an elongated molecule with dimer coiled-coil motifs (~45 nm long) that is initially largely α-helical structure, was shown to restructure with more β-sheets under tension^12, 67^. These conformational changes effectively delay the nonlinear stiffening to larger strains, allowing the structural integrity to be maintained over a wider range of strain than without such a mechanism^67^. Similarly, it has been proposed that vimentin IF could function as a flexible part of the cytoskeleton that strain hardens at very large strains as a sort of cell safety belt^8, 68^.

Dynamic assembly and maturation of vimentin intermediate filaments with strain stiffening properties as well as high compliance play a critical role in regulating cellular mechanics on different mechanical substrates^69^. Increased substrate stiffness has been correlated with increased cellular elasticity and formation of a stiff and dense network of vimentin resisting viscous flow of the cytoplasm as well as increasing the fraction of soluble versus insoluble (polymerized) vimentin^69–71^. Separately, kinase-dependent, site-specific phosphorylation of vimentin has been shown to regulate the equilibrium between vimentin assembled into filaments and disassembled subunits^72^.

Taken together, it seems that rearrangement of the IF structure in cells may play a role in mechanosensing. Early work by Fudge et al. on hagfish slime led to the initial suggestion that the formation of β-sheets within IFs might act as a sensor for local tension. Such a conformational change in response to load was proposed to possibly start a binary signaling cascade for cytoskeletal repair or apoptosis^27^. In other work by Swift and colleagues, mechanical tension, and the resulting conformational changes to the IF protein lamin-A, have been shown to inhibit phosphorylation of the lamin-A ^38, 73^. Considering the structural similarities between lamin-A and vimentin into account (both IF proteins consisting of large, α-helical coiled-coil domains), we suggest that a similar mechanism could take place within the cytoskeletal vimentin IF network. Our results on vimentin phosphorylation (**Figure 7**) agree with published work^9, 74^. They show that structural rearrangement under tension correlates with reduced phosphorylation and increased IF assembly, compared to vimentin in a reduced tension state with a more native structure. This supports the “use-it-or-lose-it” module proposed by Dingal and Discher for filamentous coiled-coil proteins^75^ since the “used” (load-bearing) vimentin is spared while the “unused” (non-load-bearing) vimentin can be depolymerized and recycled.

## Conclusion

The impact of cellular tension on the secondary structure of vimentin IFs within cells was investigated by quantitative, protein-specific BCARS vibrational microscopy. We used isotope-substitution combined with multivariate data analysis to isolate the unique isotope-substituted spectral fingerprint of deuterated vimentin from the background of cellular protein signals and accurately reconstructed the vibrational spectrum of intracellular, deuterated vimentin. These spectra provided previously unavailable information on the secondary structure of vimentin within cells. By changing the intracellular tension either pharmacologically or by growing cells on soft substrates, we found that “natural” cellular tension resulted in structural transitions of intracellular vimentin from the native, dominantly α-helical confirmation into a structure that contains more β-sheets. Our findings show that a helix-to-sheet transition of IFs occurs within cells and such a structural transition could allow vimentin to act as a local force sensor within cells.

## Acknowledgments

We are very grateful to Johannes Hunger for helpful and critical discussions. We thank Stefanie Pannwitt and Margareta Trefz for help with protein production and characterization. Sabine Pütz provided invaluable support with cell culture and Mischa D. Schwendy with confocal microscopy. Marc-Jan van Zadel and Florian Gericke provided excellent technical support for setup construction. We thank Ina Schäfer from the Core Facility Flow Cytometry, Institute for Molecular Biology Mainz, for sorting of GFP-transfected cells.

## Methods

### Protein expression and purification

A plasmid coding for human vimentin containing a C-terminal His-tag (EX-D0114-B31, Tebu-bio, Germany) was transformed into *E. coli* BL21-Gold cells. To produce deuterated vimentin, deuterated sodium acetate-d3 (Sigma-Aldrich) was added as the only carbon source to M9 minimal medium. For the production of non-deuterated vimentin, *E. coli* cells were grown in LB medium.

Bacteria were pre-cultured overnight in ampicillin (100 μg/ml) containing LB-Medium and were washed several times in M9 medium the next day. Subsequently, 1 L medium was inoculated 1:40 using the pre-culture. The growth medium contained 4 g/L deuterated sodium acetate in M9 minimal medium and ampicillin (100 μg/ml). When an optical density (OD) at 600 nm of 0.8 was reached, protein expression was induced by addition of 500 μM Isopropyl-β-D-thiogalactopyranoside (IPTG) followed by incubation at 37°C overnight. Cells were harvested the next day by centrifugation (6000 g, 10 min, 4 °C). The pellet resulting from 1 L expression culture was resuspended in 40 mL resuspension buffer (50 mM phosphate, 300 mM NaCl, 10% glycerol, pH 8.0). Cells were disrupted on ice by sonication (Omni Sonic Ruptor 400, Omni) and inclusion bodies were separated via centrifugation (30 min, 6000 g, 4 °C). Next, the inclusion bodies were solubilized in lysis buffer (100 mM NaH_2_PO_4_, 10 mM Tris-HCl, 8 M urea, 150 mM NaCl, pH = 8.0) for 3 h at room temperature. Cell debris was removed by centrifugation (15 min, 6000 g, 20 °C) and the supernatant was incubated overnight with 1 mL Ni-NTA agarose suspension at 4 °C. The solution was centrifuged (3 min, 800 rpm, 20 °C) and the pellet was resuspended in 5 ml of the supernatant and loaded onto an empty column. The Ni-NTA-agarose was then washed four times using 5 mL washing buffer (100 mM NaH_2_PO_4_, 10 mM Tris-HCl, 8 M urea, 150 mM NaCl, pH 8.0) with increasing imidazole concentrations (0, 5, 10, 15 mM). Vimentin was eluted twice from the column with 1 mL 400 mM imidazole in 100 mM NaH_2_PO_4_, 10 mM Tris-HCl, 8 M urea, 150 mM NaCl, pH 8.0. The fraction containing the target protein was dialyzed stepwise against a series of buffers with decreasing urea concentrations (6, 4, 2 M urea in 5 mM Tris-HCl, 1 mM DTT, pH 8.4) at 4 °C for 1 h each, followed by an additional dialysis against fresh buffer (5 mM Tris-HCl, 1 mM DTT, pH = 8.4) overnight at 4 °C. The final solution was analyzed by SDS-PAGE and Coomassie-staining, showing only a protein band with an apparent molecular mass of about 55 kDa, as indicated by a protein ladder (PageRuler Prestained, Thermo Scientific).

### *In vitro* stretching of IF

Vimentin IFs were polymerized *in vitro* by adding NaCl to a final concentration of 170 mM to a 1.2 mg/ml vimentin monomer solution^76^. Filaments formed with a 2 h incubation of the solution at 37 °C. The IF containing solution was pipetted onto a coverslip, and filaments (and bundles) were allowed to settle onto the glass surface (see also **Figure S21**). Next, attached filaments were deformed by gently pulling with a glass capillary (Femtotip, Eppendorf) mounted to a micromanipulator (Injectman II, Eppendorf).

### Cell culture

HeLa-GFPvim cells, expressing vimentin with a GFP tag were produced by lentiviral transfection (LentiBrite GFP-Vimentin Lentiviral Biosensor, Merck Millipore) of HeLa cells. To establish a strain with sufficient high (>60% of cells) expression rate of GFP, the cells were sorted by fluorescence-activated cell sorting.

For all microscopy imaging experiments, HeLa-GFPvim were grown in collagen-coated glass bottom dishes (MatTek) and in culture medium (Dulbecco’s Modified Eagle’s Medium + 10% fetal calf serum, Dulbecco) containing 100 U/ml Penicillin and 100 μg/ml Streptomycin (Gibco).

### Disruption of native IF cytoskeleton

To achieve incorporation of deuterated vimentin into the cellular vimentin network, the native IF network was depolymerized by cycloheximide prior to microinjecting. This treatment is known to alter the IF network while actin stress fibers and microtubules remain unaffected^48^. Cycloheximide (ready-made solution in DMSO, Sigma Aldrich) was diluted in DMEM containing 10% FCS to a final concentration of 10μg/ml, and the cycloheximide solution was left on the cells for 6 h at 37°C, 5% CO_2_ and 95% relative humidity. The IF depolymerization was reversible within 12 h after changing back to normal conditions. Successful disruption was checked by fluorescence microscopy (see SI **S3**).

### D-Vim microinjection

For microinjection, the cell medium was changed to 4°C cold Leibovitz’s L-15 CO_2_ independent medium (Gibco). Prior to microinjection, concentrated vimentin monomer solution (4 mg/ml) was centrifuged at 13000 rpm for 5 min to prevent capillary clogging from protein aggregates. The supernatant was used for microinjection into adherent HeLa-GFPvim cells with a microinjection system (Injectman II and Femtojet, Eppendorf), and injection parameters were 100 Pa injection pressure, 0.5 s injection time and 50 Pa compensation pressure. After injection, the medium was changed back to normal culture medium and cells were left to incubate for 12 h to allow cells recover and reform the IF network.

### Alteration of cellular tension

The cytoskeleton tension was changed either by the chemical agent blebbistatin or physical modification of the cell substrate. Blebbistatin (Abcam), a small molecule known to inhibit myosin-II in actin stress fibers^52^, was added to culture medium containing 10% FCS to a final concentration of 50 μM. For chemically induced relaxation, HeLa-GFPvim cells containing isotopically labeled D-Vim and grown on rigid collagen-coated glass substrate were incubated in the blebbistatin-containing medium for 8 h.

In a second series of experiments, cells were grown during all treatment steps (depolymerization, microinjection, repolymerization) on substrates of low stiffness – collagen-coated softwell hydrogel of 0.1 kPa elastic modulus (Softview, Matrigen) – instead of glass. This was done to maintain a constant (collagen) surface functionalization to the rigid glass bottom dishes used in other experiments.

In each experiment, cells were prepared for BCARS by washing the cells with PBS followed by fixation for 30 min (4% paraformaldehyde in PBS, pH 7.4). The cell samples were stored in PBS at 4°C and used for BCARS experiments within 3 days. Morphological changes of the IF network were visualized by confocal microscopy (see Supplementary Methods for details).

### BCARS Microspectroscopy

Fixed cell samples were analyzed with a home-built broadband anti-Stokes Raman Scattering (BCARS) spectroscopy setup that has been extensively described elsewhere^77^. A brief description of the experimental setup can be found in the supplementary information (see SI **Figure S20**). Using this setup, a region of interest, usually covering a single cell, was raster scanned and a BCARS spectrum was recorded at each pixel.

As the acquired BCARS signal consists of both nonresonant and resonant contributions from the sample and the surrounding medium, a phase retrieval procedure was performed to isolate Raman-like spectra for quantitative analysis^78^. The method described by Liu *et al.* using a time-domain Kramers-Kronig transform with a causality constraint was employed to produce the resonant contribution from each BCARS spectrum with a nonresonant spectrum acquired from the nearby medium^79, 80^. However, due to this imperfect reference, the slowly altering error from the phase had to be corrected. This was achieved by subtracting a spectral baseline which was calculated by an iterative fit with a fifth-order polynomial^81^. The entire spectral processing to obtain Raman-like spectra was performed by home-written scripts in Igor Pro 6.37 (WaveMetrics). Representative hyperspectral data can be found in the Supplementary Information (see SI **Figure S7** and **S8**).

### Separation of spectral contribution from isotopically labeled protein

To separate the Raman signal of deuterated vimentin IF from all other protein contribution, we employed multivariate curve resolution using an alternating least squares algorithm (MCR-ALS)^82^ on every BCARS dataset covering a single cell. The processing was done with the MCR-ALS GUI 2.0 toolbox for Matlab (R2015, MathWorks) by Tauler and coworkers^83^. A similar approach was established to isolate the spectrum of a deuterated peptide within cells by our group in an earlier study^53^.

In a first step, every RL-dataset was filtered against a threshold set for the CH band intensity (2935 cm^−1^) to exclude pixel outside the target cell. From the remaining pixels, an average spectrum was calculated. Next, the spectral data was reduced further to spectra showing at least some traces of D-Vim as evaluated by the presence of the CD_2_ vibration. As our interest here only was on the secondary structure and the spectral labeling in the CD region, the spectral range was narrowed the following regions: 1400 cm^−1^ - 1780 cm^−1^ and 2130 cm^−1^ - 2400 cm^−1^. Thereby, extraneous spectral features as well as intrinsic noise in the quiescent region were filtered out and the prepared spectra were bundled into a single data matrix per cell. Next, MCR-ALS was used to find the optimal pure spectral components and respective loadings to represent the entire dataset. A maximum of three spectral components (for cell background, D-Vim and non-random artifacts) was chosen together with constraints for non-negativity of loadings and spectra. The iterative process was stopped either when convergence was achieved or after 50 steps. From the calculated component spectra, the D-Vim component was selected as that having the highest signal intensity in the CD part of the spectrum. From those, an average spectrum was calculated and analyzed further to judge the secondary structural content.

### Calculation of secondary structure

Raman spectroscopy is sensitive in several vibrational modes to secondary structure motifs of molecules present in the sample volume^40, 42^. This dependency was used to calculate the structural composition from the given average RL spectra as well as from component spectra obtained by MCR analysis as described above. The Amide I band (1570 – 1730 cm^−1^) was decomposed via least square fitting with a Levenberg-Marquardt algorithm in Matlab (Mathworks). Thereby, the Raman modes assigned to different structural motifs with known peak position, were extracted. From the resulting peak areas, the overall structural composition was calculated for the given spectrum. As shown by Berjot et al., the Amide I band can be decomposed into two peaks for α-helix at 1635 and 1647 cm^−1^, a peak at 1660 cm^−1^ for random coils, 1667 cm^−1^ for β-sheets and 1693 cm^−1^ for β-turn. Ring modes from tyrosine, tryptophan and phenylalanine were included with two minor peaks at 1600 cm^−1^ and 1612 cm^−140^. The peaks were assumed to have Lorentzian line shape, defined by a linewidth and center frequency both being allowed floating within a defined range. The major fitting parameter, the peak amplitude, was constrained to only positive values.

### Vimentin-phosphorylation on different mechanical substrate

GFP-Vimentin expressing HeLa cells were cultured on sterile glass slide (10 x 10 mm) and soft collagen hydrogel (3 mg/ml, Gibco) in 24 well plate as described previously^84^. After 24 h of culture, cells were fixed using 4% formaldehyde solution, permeabilized using 0.1% Triton solution in PBS for 15 min and blocked with 1% BSA (Sigma) for 2 hr. Cells were incubated with primary mouse anti-Vimentin (phosphor-Ser55)(pSer55)(1 μg/ml; Enzo Life Sciences) antibody in 1% BSA over overnight at 4 °C. Later cells were rinsed in PBS several time and incubated with secondary antibody Goat anti-Mouse IgG (Alexa Fluor 568) in 1% BSA for 1 hour at room temperature and rinsed multiple time with PBS. Finally, antibody stained cells were imaged with Olympus Fluoview FV3000 confocal microscope with 40X objective having 0.6 NA.

