## Supporting Information for "Tension causes structural unfolding of intracellular intermediate filaments"

**Supplementary Methods**

Confocal microscopy

Confocal microscopy was used to visualize the spatial distribution of vimentin IF in HeLa-GFPvim cells under different conditions. Measurements were performed on a TCS SP5 confocal microscope (Leica, Germany) with laser excitation at 405 nm (GFP) and a 63x, 1.2 NA water immersion objective lens. The image contrast was adjusted and a smoothed background was removed in ImageJ.

Vimentin at a concentration of 0.2 mg/ml were allow to assemble at 37 °C by adding 5 mM sodium phosphate buffer containing 150 mM NaCl. After 30 min of assembly, vimentin was labelled using Alexa Fluor 488 anti-vimentin antibody (0.5 µg/ml) (BioLegend, USA). For microscopic screening of vimentin filaments, 50 µl of labelled vimentin allowed to adsorb on to cleaned cover slide for 15 min followed by gentle rinsing with distilled water. Fluorescence imaging was performed using confocal laser scanning microscope (Olympus Fluoview FV3000) with 30X, 1.05 NA silicon oil immersion objective.

Widefield fluorescence microscopy

To determine the intracellular distribution if native IF under different treatments as well as microinjected vimentin, widefield fluorescence microscopy was used. Measurements were done on an Olympus Ti-81 microscope with 20x or 100x magnification with appropriate filters.

Circular dichroism spectroscopy

Secondary structure of vimentin monomers was analyzed with circular dichroism spectroscopy in the range from 190 to 260 nm in 1 nm steps. Experiments were performed on a J-815 spectrometer (JASCO Inc, Easton, USA) under nitrogen atmosphere and by using quartz cuvettes of 1 mm path length. Protein was dissolved in phosphate buffer at pH 7.4 with final concentration of 2.5µM. For each sample, 3 spectra were taken and averaged.

CARS microspectroscopy

Intracellular protein structure was analyzed by broadband CARS microspectroscopy setup as depicted in **Figure** **S20**. A commercial ns-pulsed laser source (Leukos-CARS, Leukos) provides a spectrally narrow pump/probe beam (λ = 1064 nm) from which ~ 50% is coupled into a photonic crystal fiber to obtain a spectrally broad Stokes beam (λ = 400-2400 nm). To match both Stokes and pump/probe beam pulses in the sample focus temporally, the later one is guided through a delay line of several meters length. The spectral range of the Stokes beam is cut by a longpass filter and a Glan-Thompson polarizer to a bandwidth from 1100-1600 nm. The final spectral power density of the Stokes beam is more than 100 µW nm^-1^. Both Stokes and pump/probe beam are combined by a dichroic mirror and directed to an inverted microscope (Eclipse Ti-U, Nikon) where they are focused into the sample with an objective lens (100x, NA:0.85, Zeiss) as shown in **Figure** **S20 B**. The total average laser power within the focal volume is 30 mW. To allow hyperspectral imaging, the sample is mounted on a xyz piezo stage (Nano-PDQ 375 HS, Mad City Labs) allowing sub-micrometer step size. The CARS signal is collected in forward direction with a second objective lens (10x, NA: 0.25, Zeiss) and send to a spectrometer. To block the intense pump/probe and Stokes beams, a notch-filter (NF03-532/1064E-25, Semrock) and a short-pass filter (FES1000, Thorlabs) are placed before the entrance slit. The CARS signal is analyzed by a Czerny-Turner spectrograph (Shamrock 303i, Andor) with attached cooled CCD camera (Newport DU920P-BR-DD, Andor) resulting in a spectral range from 500 cm^-1^ to 4000 cm^-1^ with a spectral pitch of ~ 4 cm^-1^. All components in the setup are controlled with software written in LabView (National Instruments). Further details of this BCARS experiment can be found elsewhere ^1, 2^.

Cell treatment with latrunculin-A

Cells pre-injected with D-Vim were treated with latrunculin-A to depolymerize actin filaments reduce cell tension. Latrunculin-A (Abcam) was incubated with cells at a final concentration of 5 µM for 1 h in standard medium. Following this incubation period, cells were treated identical as described in the Methods (Alteration of cell tension) for BCARS imaging.

**Supplementary Data**

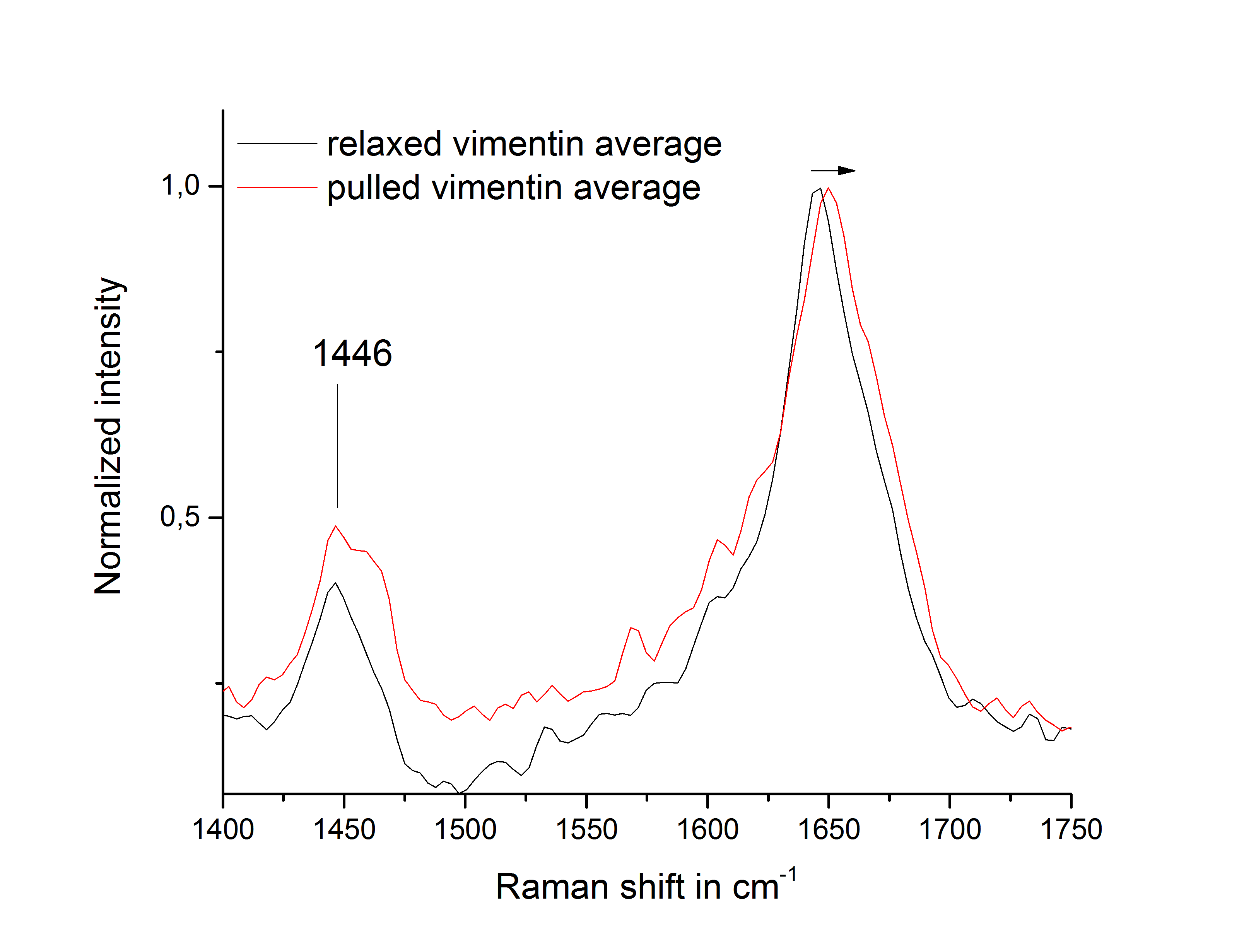

**Figure S1:** Pulling *in vitro* vimentin filament bundles showed spectral shifts in the Amide I region. A significant difference in the Amide I peak position, with a shift to higher wavenumbers for the mechanically stressed samples, was observed. The spectra from relaxed vimentin filament bundles and vimentin bundles pulled with a micromanipulator tip (to more than 100% strain) were averaged over 3 samples per group and averaged 10 spectra from each sample. For the relaxed vimentin an average peak center of 1645 ±1.5 cm^-1^ was found while for the pulled vimentin samples the center was 1651 ±1.0 cm^-1^. A two-sample t-test assuming unequal variance resulted in a significant difference of the peak center frequencies.

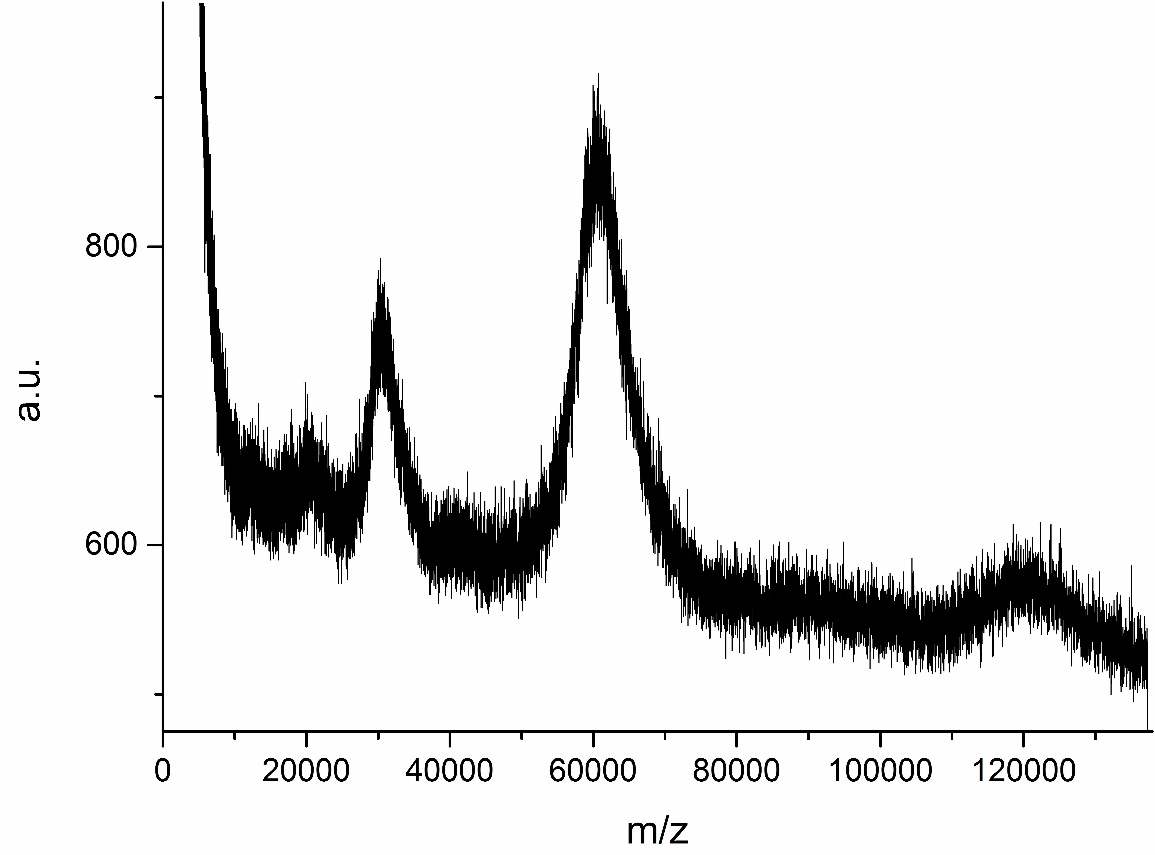

**Figure S2:** Maldi-TOF spectrum of recombinantly produced deuterated Vimentin showed maxima at 60654.45 m/z and 30243.68 m/z, indicating almost full isotopic exchange after production.

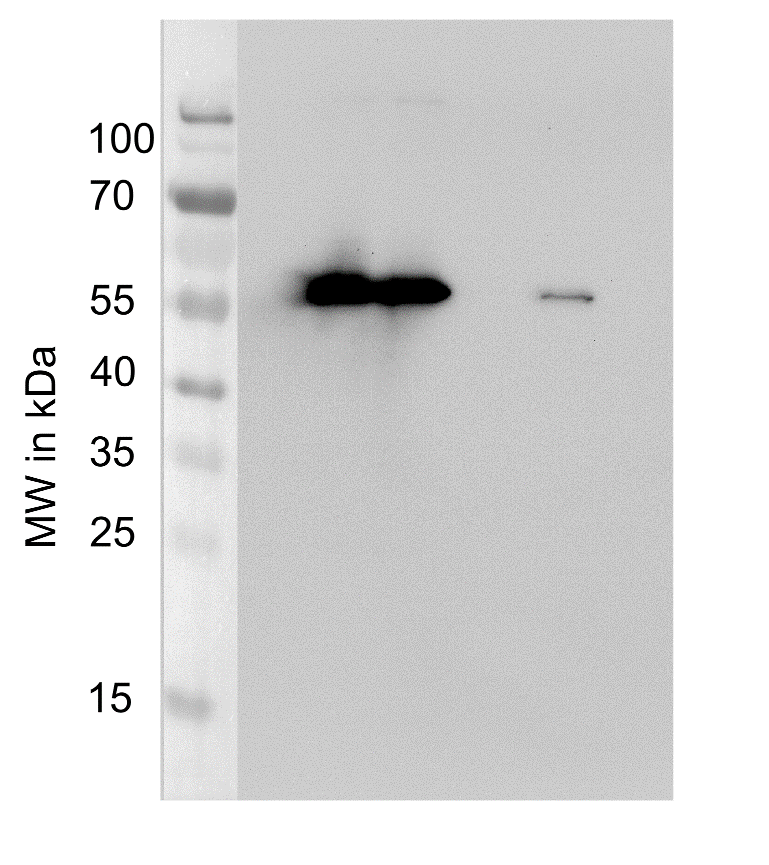

**Figure S3:** Western Blot showed vimentin of high (first two lanes) and low concentration (fourth lane) by an anti-his labeling. The expected MW is about 55 kDa.

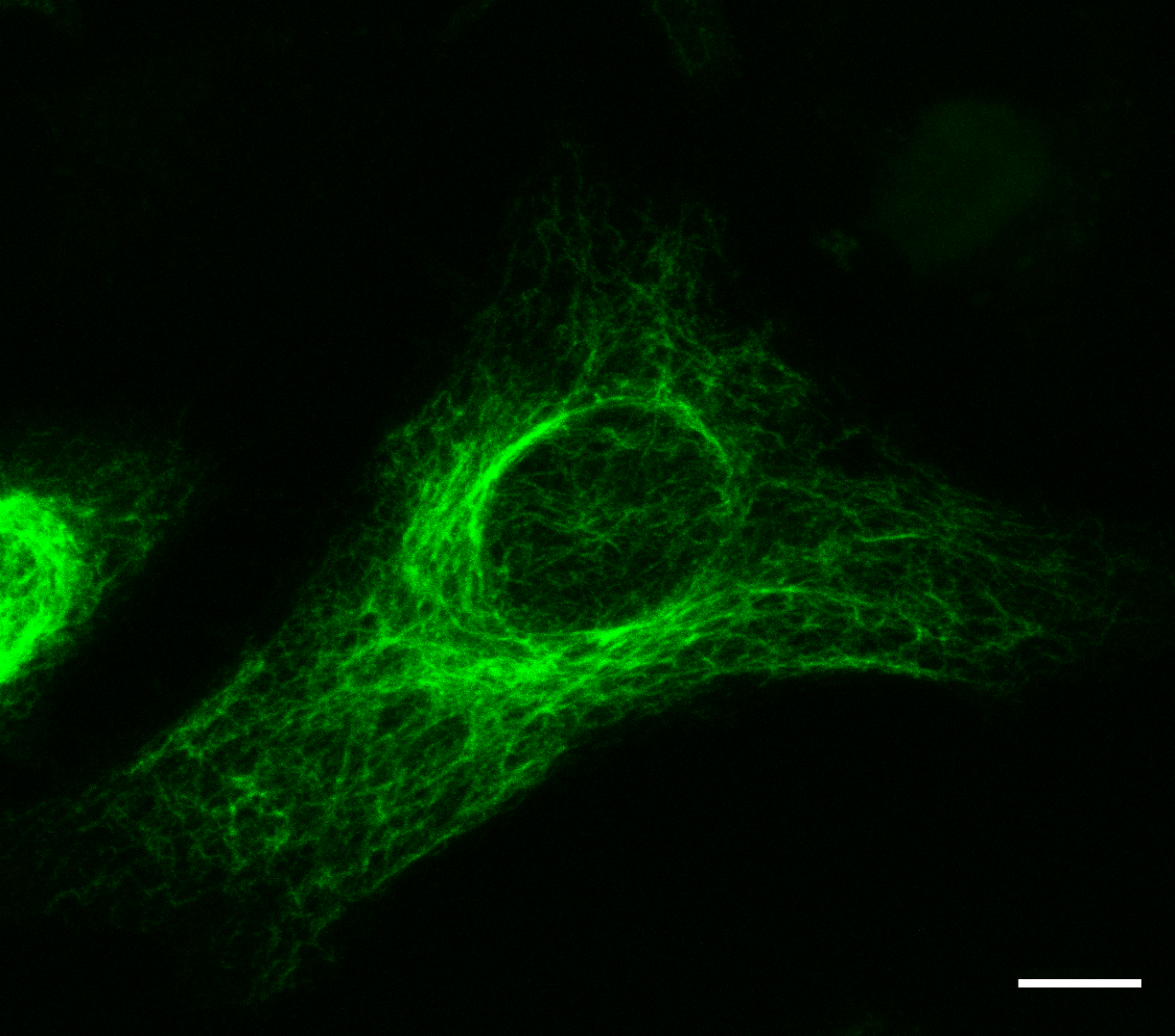

**Figure S4**: Confocal microscopy image of HeLa cells grown on collagen-coated rigid glass surface co-expressing Vimentin and GFP (strain Hela-GFPvim). The IF network was filamentous and spread out through the entire cytosol, being most dense in close proximity to the nucleus. Scale bar is 10 µm.

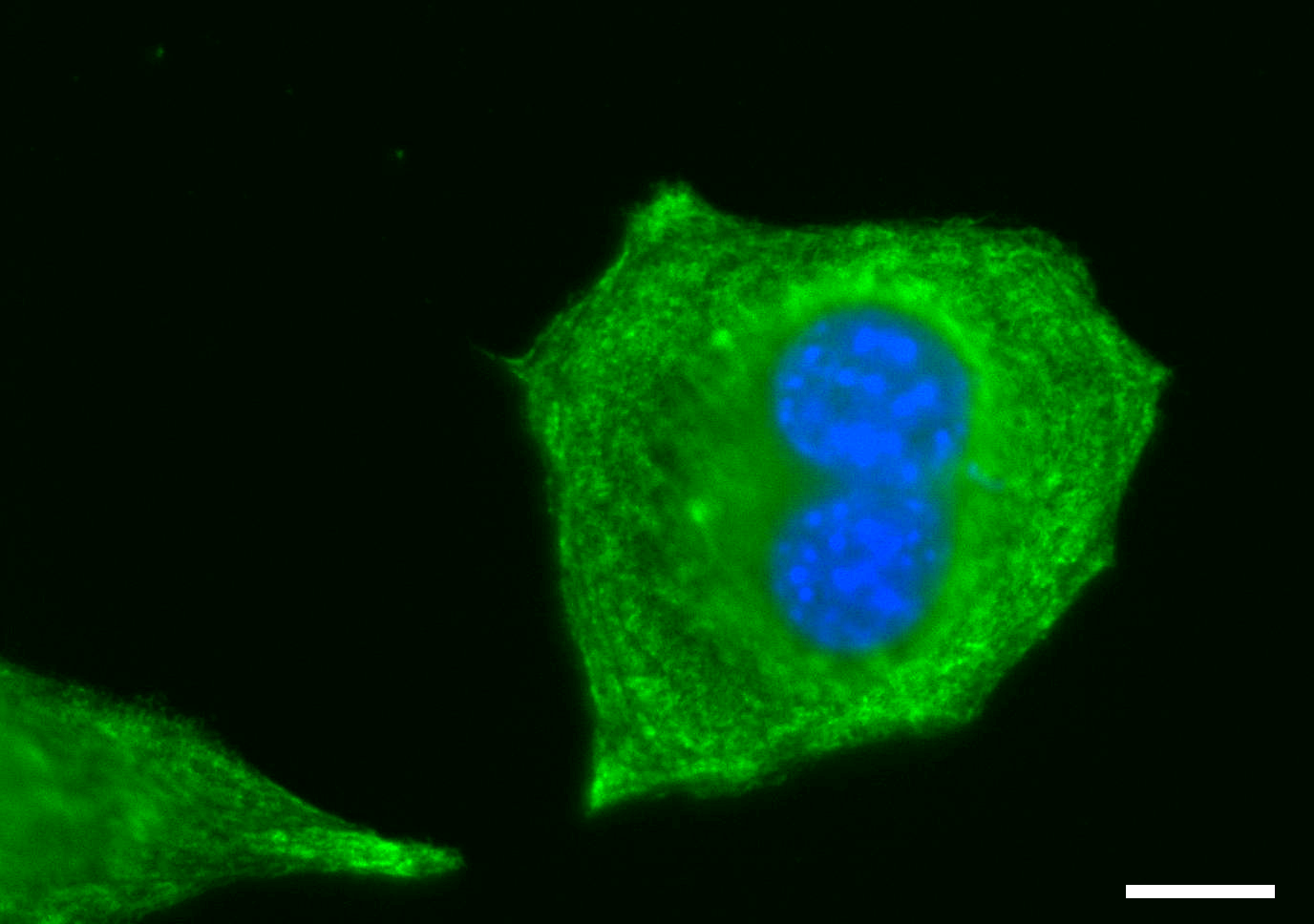

**Figure S5**: Merged widefield microscopy image showing the disassembled GFP-labelled (green channel) vimentin IF network in HeLa-GFPvim cells on petri dishes after 5 h of treatment with cycloheximide. Nuclei were labelled by DAPI in blue. Scale bar is 10 µm.

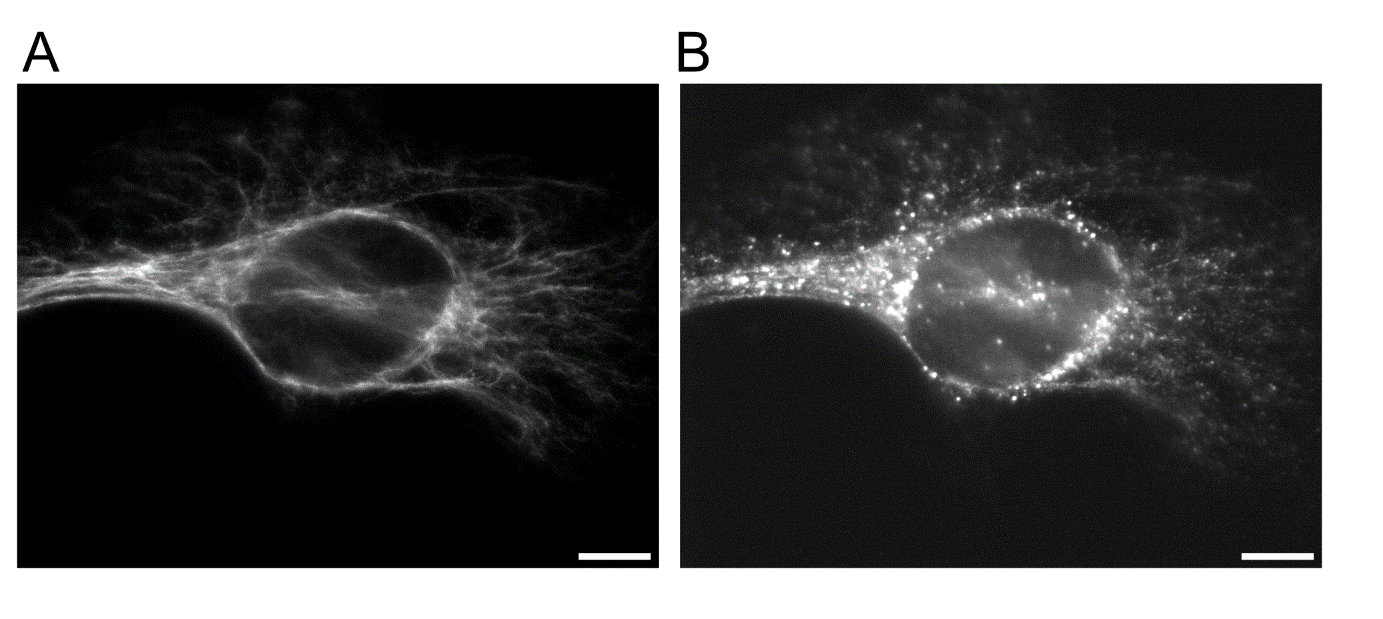

**Figure S6:** Incorporation of microinjected vimentin into the native IF network. (A) Widefield fluorescence microscopy image of GFP-labelled native IF in HeLa after ~8 h recovery from depolymerization with cycloheximide. (B) Injected, rhodamine-labelled vimentin resembled the same pattern as the intrinsic vimentin. The bright dots are caused by dye aggregates. Scale bars are 10 µm.

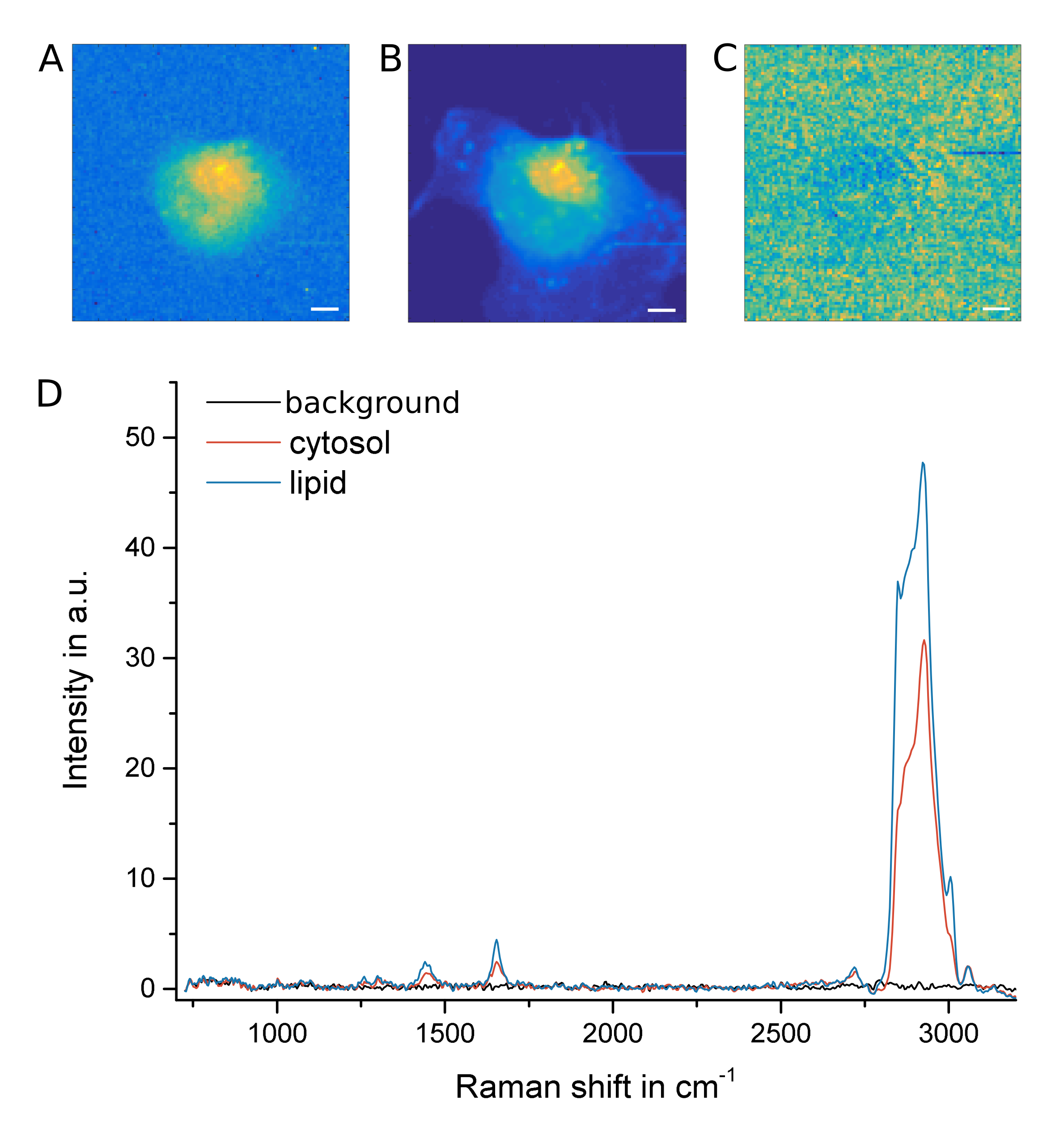

**Figure S7:** Broadband CARS (BCARS) data of a cell without the carbon-deuterium (CD) peaks from deuterated vimentin. Hyperspectral maps of A) the Amide I band, B) CH band between 2800 and 3200 cm^-1^ and C) CD band between 2100 and 2250 cm^-1^. Scale bars are 5 µm. D) Representative spectra from outside the cell (‘background’), from the cytosol (‘cytosol’) and a lipid droplet (‘lipid’). No Raman signal appeared in the silent region between 1700 and 2600 cm^-1^.

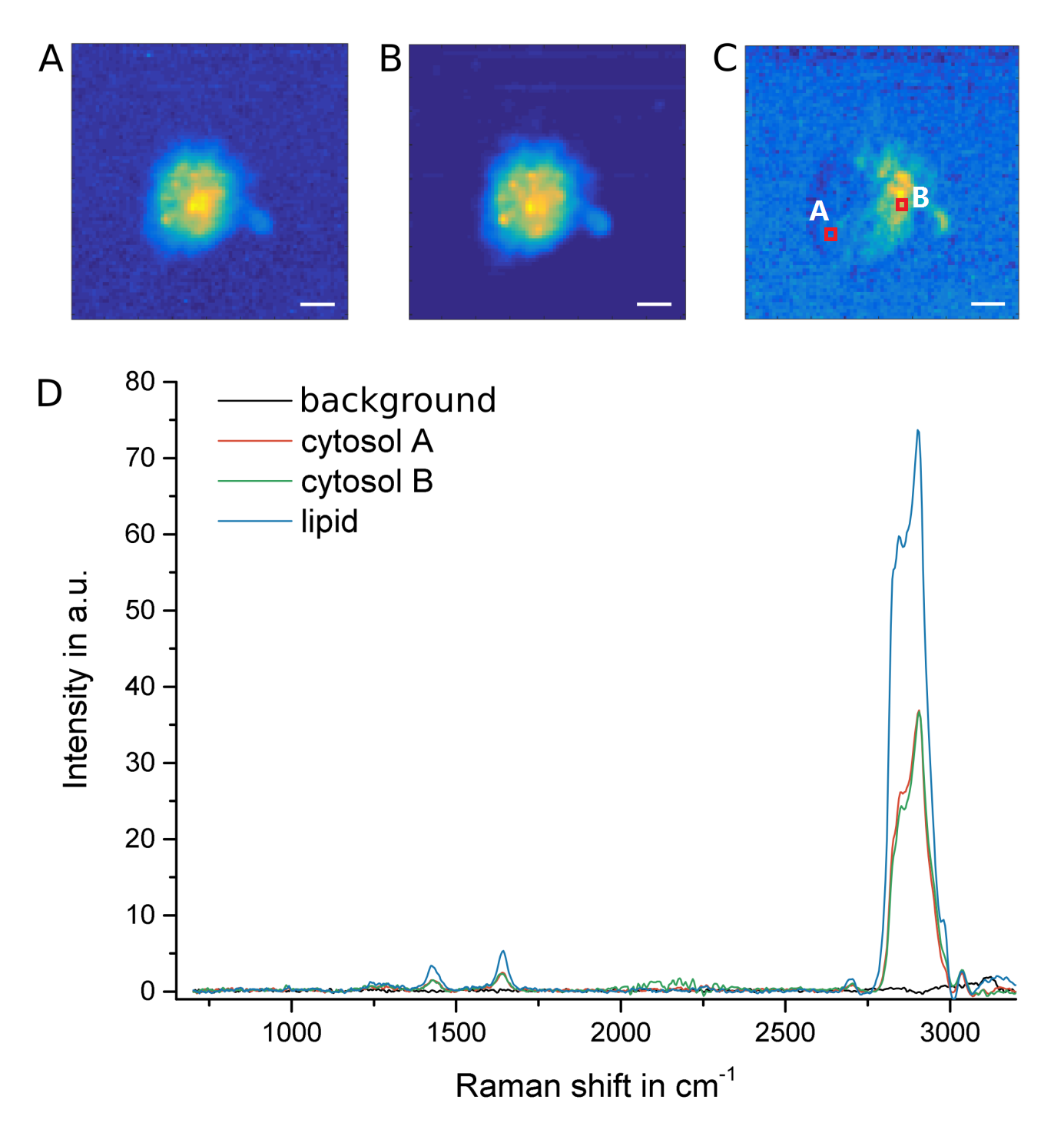

**Figure S8:** Cell containing deuterated vimentin. Hyperspectral maps of A) the Amide I band, B) CH band between 2800 and 3200 cm^-1^ and C) CD band between 2100 and 2250 cm^-1^. Scale bars are 5 µm. D) Representative spectra from outside the cell (‘background’), from two different positions in the cytosol (‘cytosol A’ and ‘cytosol B’): one without d-Vim and another rich in d-Vim, and a lipid droplet (‘lipid’). The additional signature of the CD mode appears in the silent region around 2150 cm^-1^.

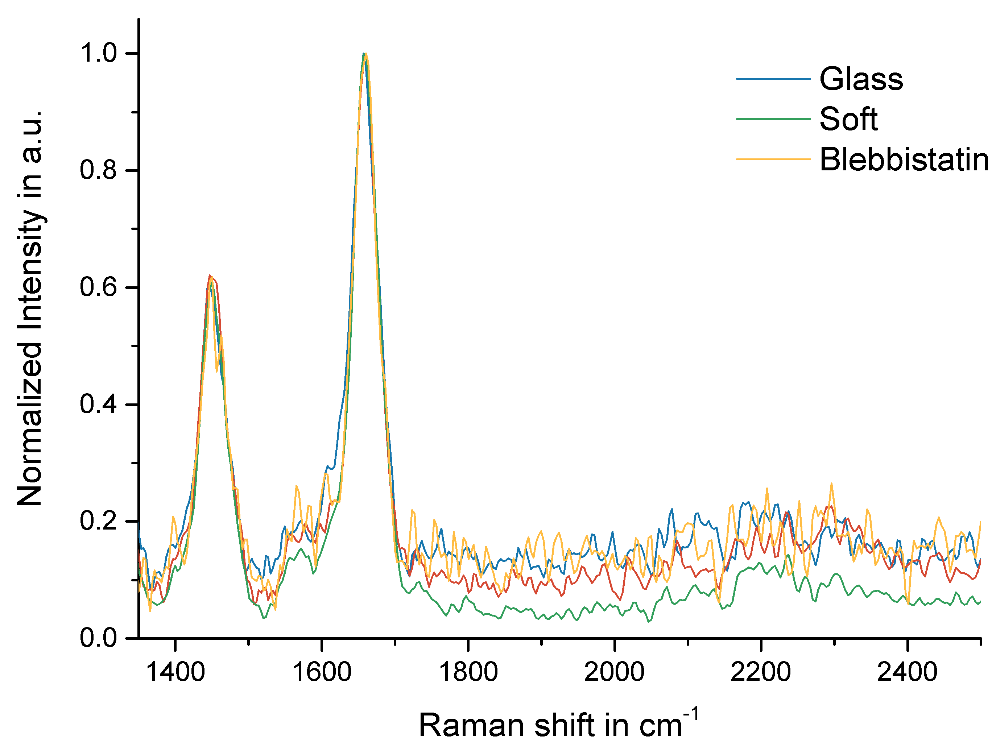

**Figure S9:** Normalized average cell spectrum for each cell group showed a constant Amide I region. The spectra per group show were calculated by pooling all spectra originating from cell-containing pixels from each cell for each group and taking the global average per group.

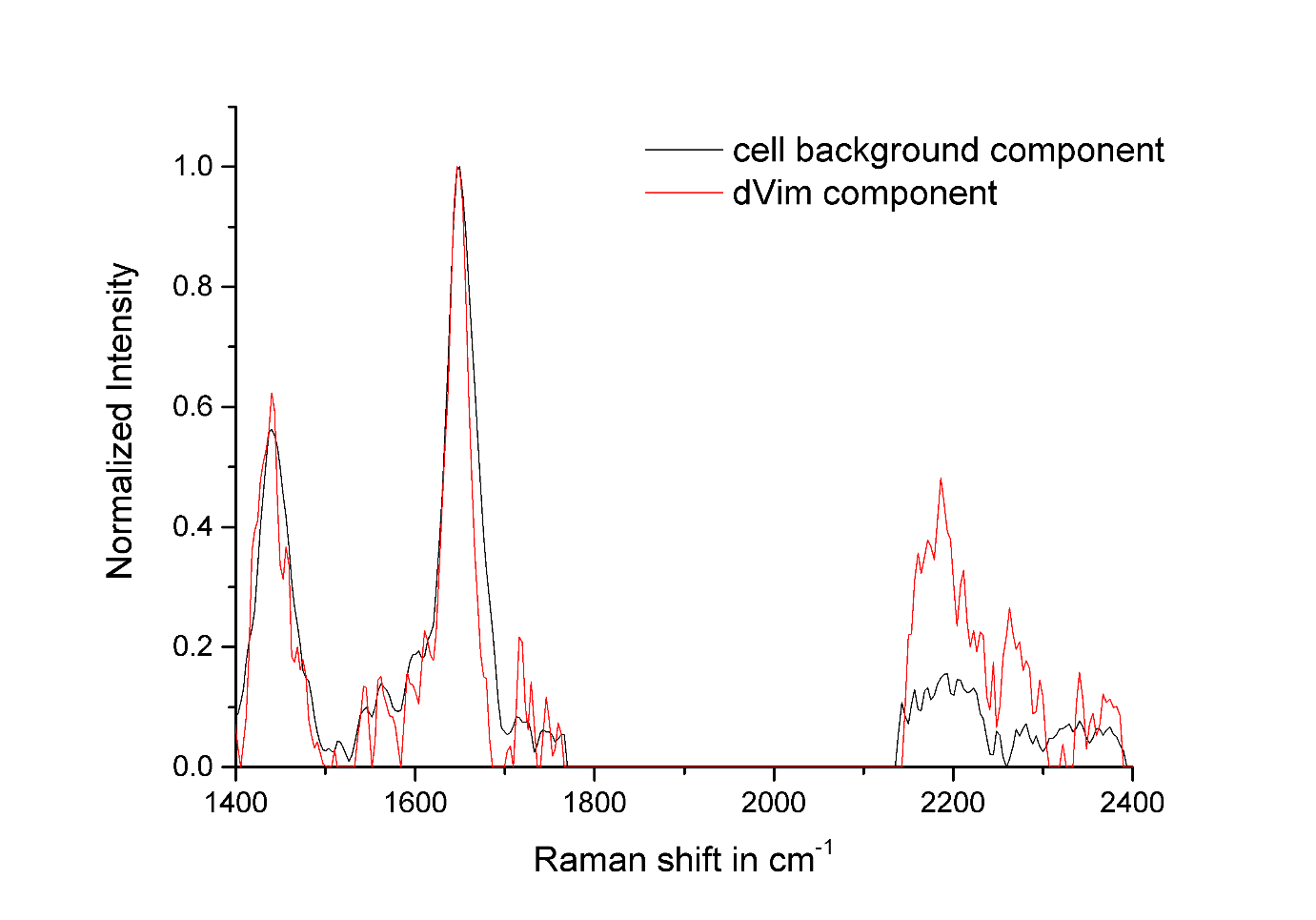

**Figure S10:** Average of the MCR components obtained from CD-containing pixels from cells treated with blebbistatin that were grown on collagen-coated glass substrates. The two components differed mostly in their intensity of the CD region with some changes in the Amide I and CH region. The component with the strongest CD region intensities was assigned to the deuterated protein while the remaining component represents any non-deuterated protein.

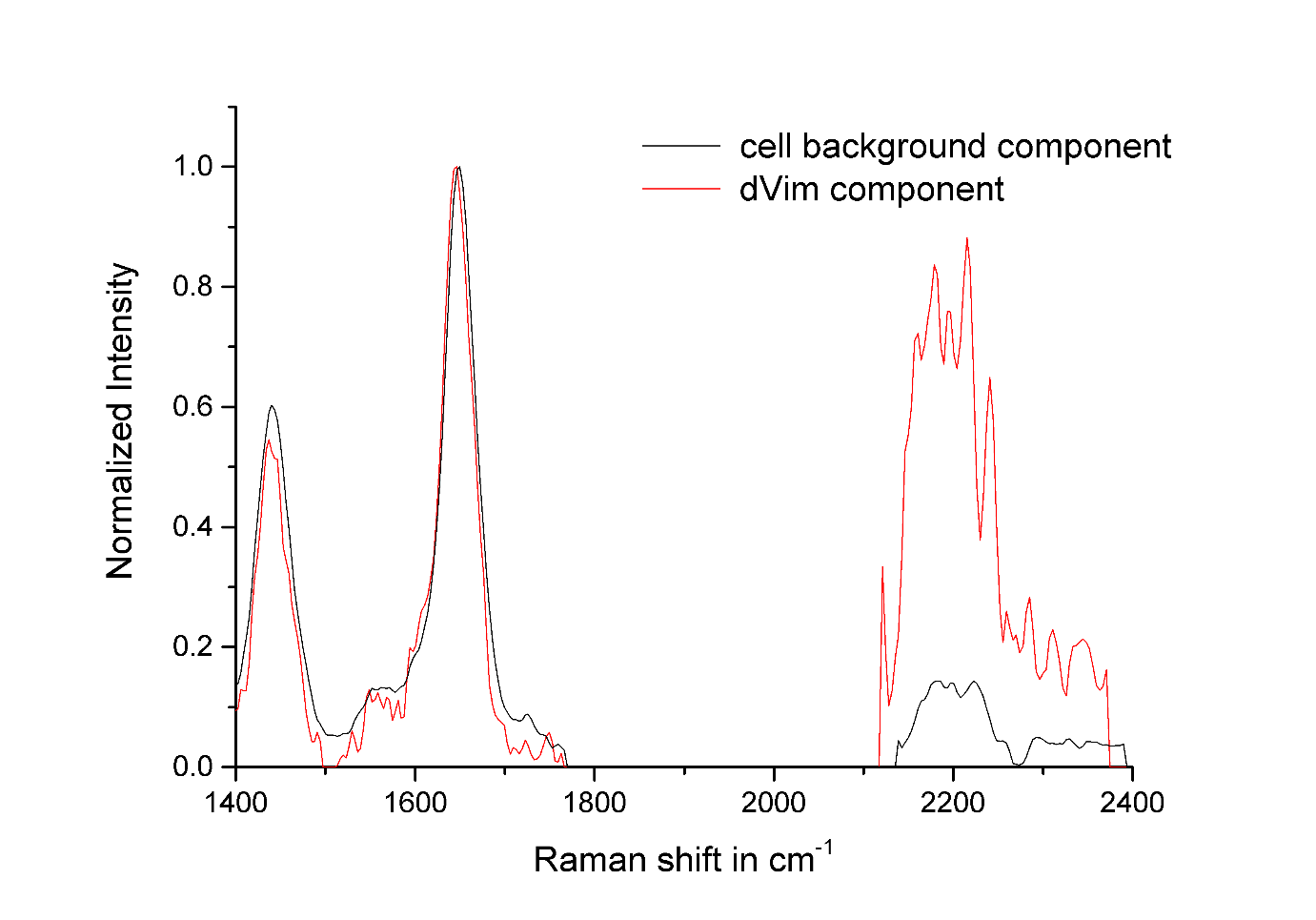

**Figure S11:** Average of the MCR components obtained from CD-containing pixels of cells that were grown on collagen-coated soft substrates. The two components differed mostly in their intensity of the CD region with minor changes in the Amide I and CH region. The component of strong CD intensities was assigned to the deuterated protein while the remaining component represents any non-deuterated protein.

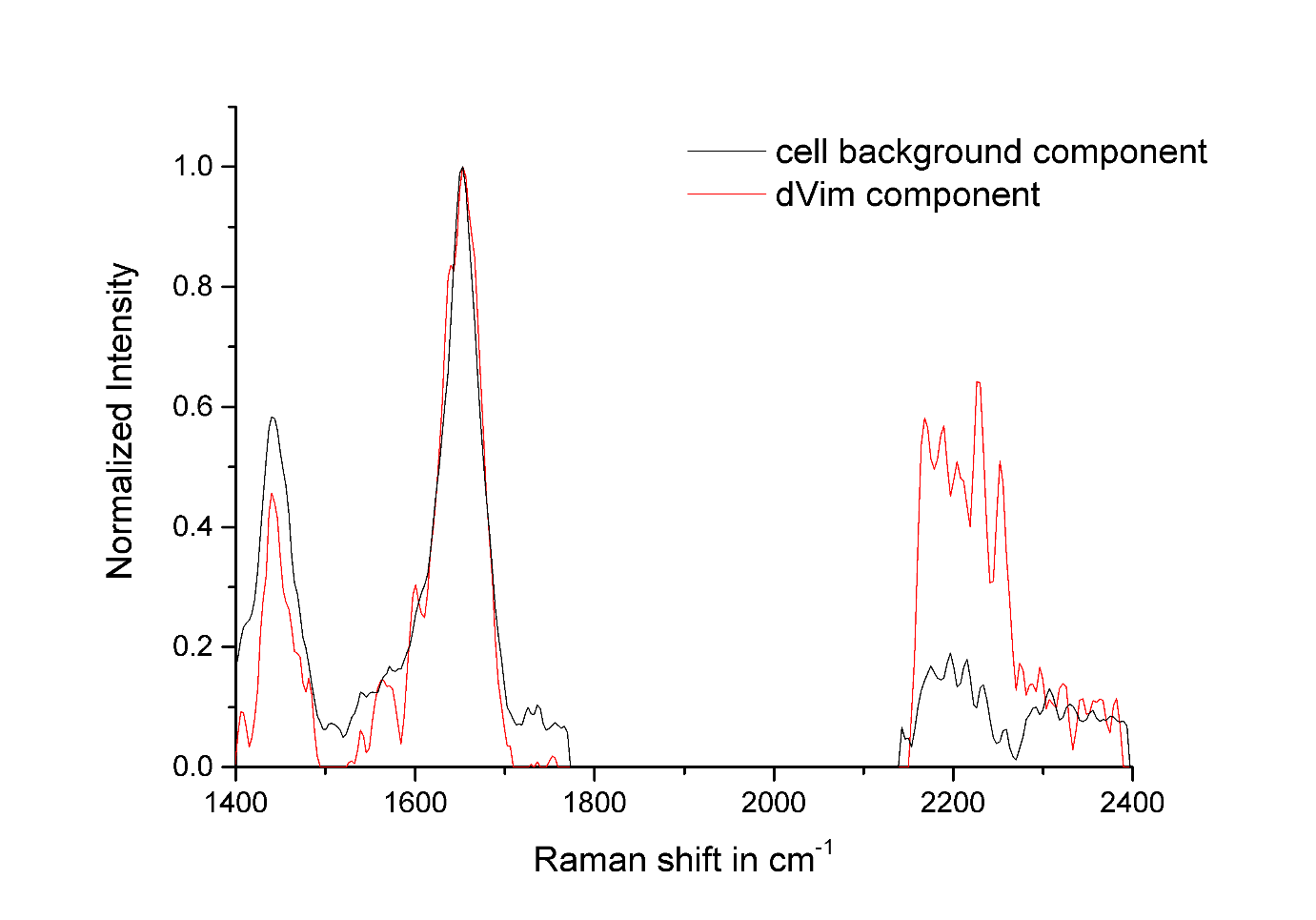

**Figure S12:** Average of the MCR components obtained from CD containing pixels of cells that were grown on collagen-coated glass substrates. The two components differed mostly in their intensity of the CD region with some changes in the Amide I and CH region. The component of strong CD intensities was assigned to the deuterated protein while the remaining component represents any non-deuterated protein.

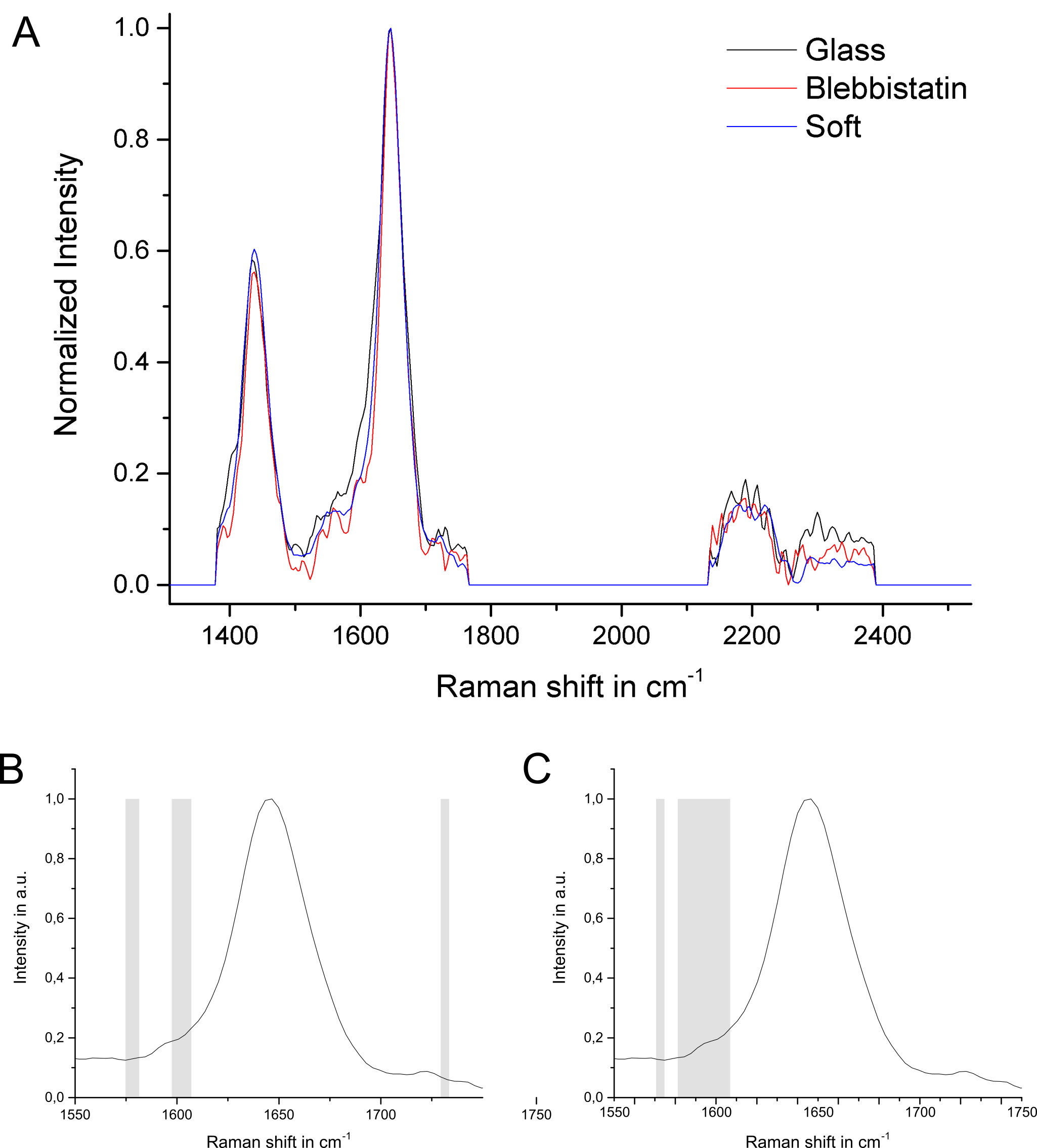

**Figure S13**: MCR component for the cellular background from different groups normalized to the Amide 1 maximum at 1652 cm^-1^. The shown spectra represent the average of all cells per group (A). Statistical differences determined by a two-sample t-test for glass and blebbistatin (B) and glass and soft substrate (C) is indicated in gray.

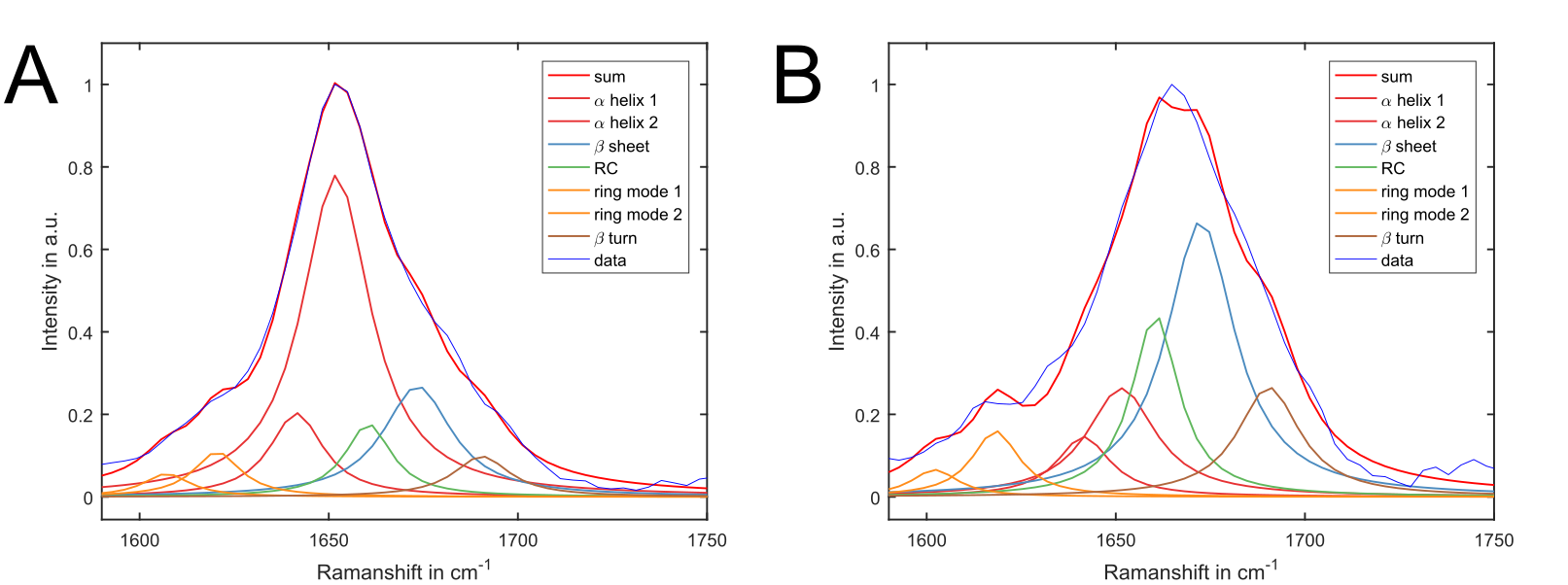

**Figure S14**: Amide fit on the average spectral data from pure vimentin (A) and as average over an entire cell (B).

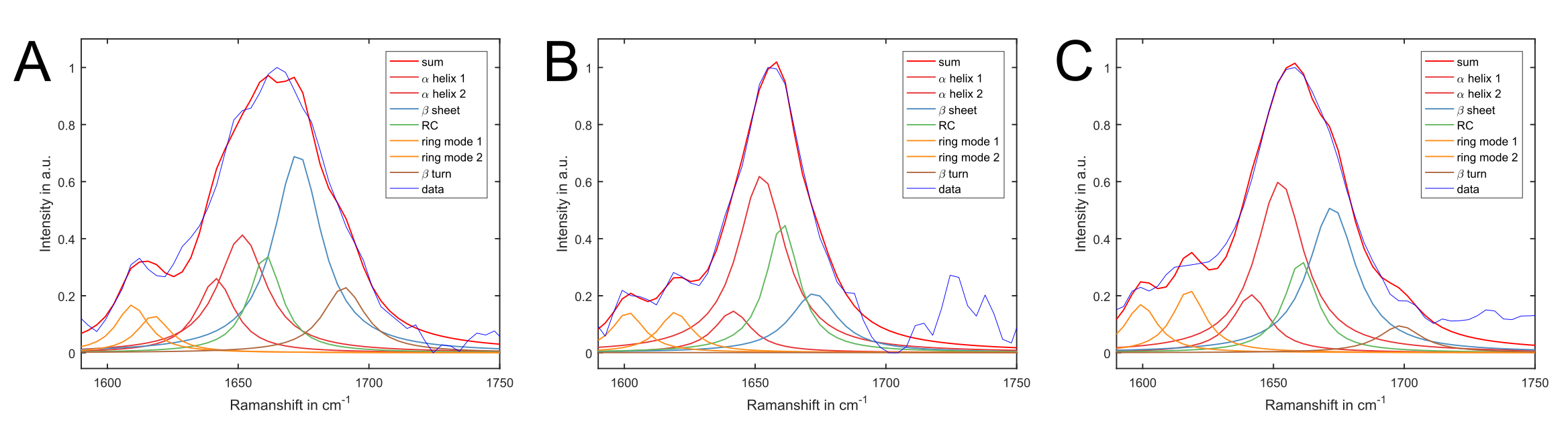

**Figure S15**: Amide fit on the average MCR components assigned to D-Vim from cells grown on glass (A), treated with blebbistatin (B) and grown on soft substrate (C).

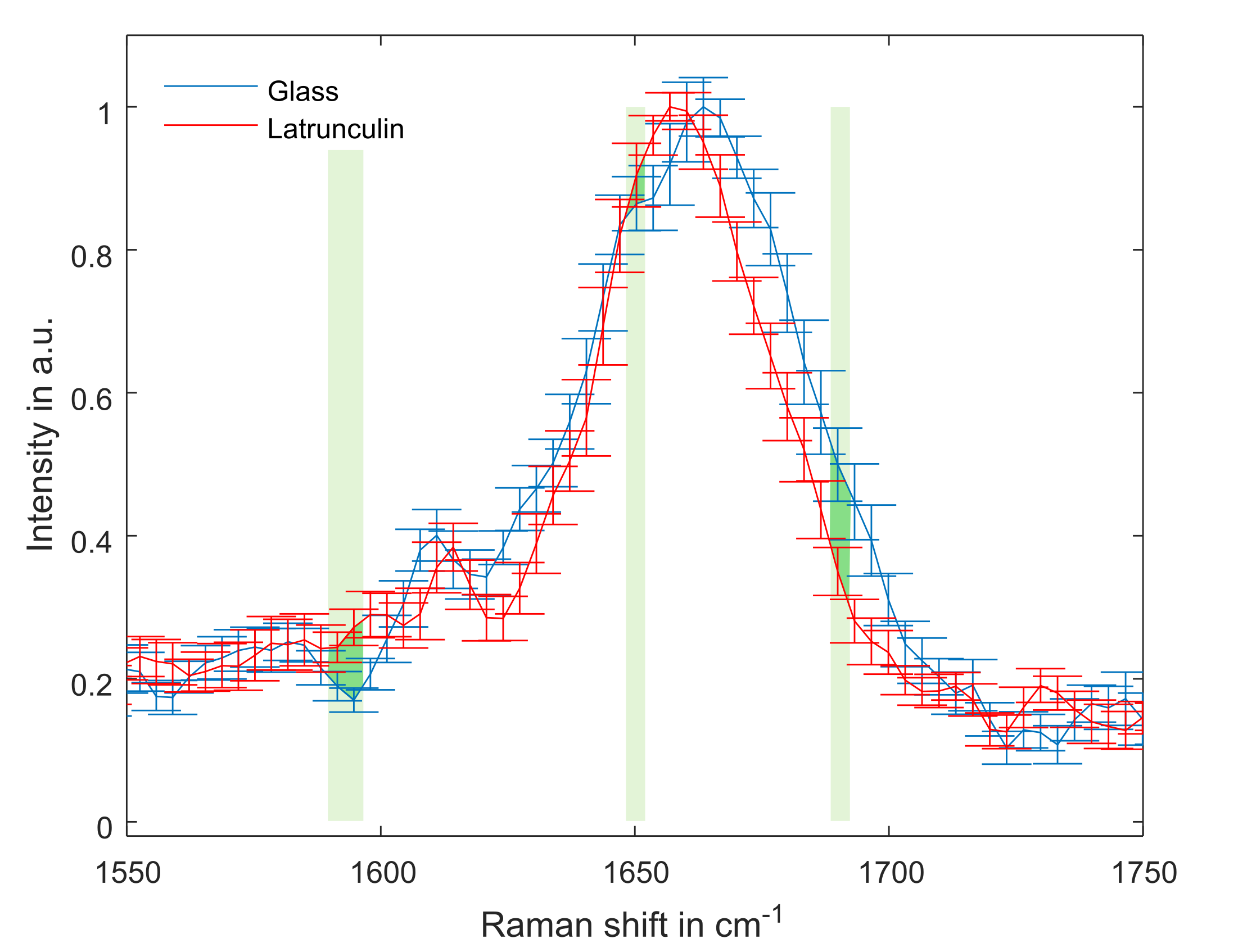

**Figure S16:** Spectral analysis of latrunculin-A treated cells compared to the control group showed changes in the Amide I region. Average component spectra of cells from collagen-coated glass substrates (n=14) and cells from the same substrate after Latrunculin-A (n=18) treatment. The SEM is shown as error bars for each wavenumber data point and significant differences are indicated by green shading.

| **Structure** | **Glass average [%]** | **Glass SEM [%]** | **Lat-A average [%]** | **Lat-A SEM [%]** |
| --- | --- | --- | --- | --- |
| α-helical | 32.5 | 3.0 | 39.0 | 4.7 |
| β-sheet | 35.7 | 2.1 | 31.6 | 2.8 |
| RC | 31.8 | 3.2 | 29.5 | 3.0 |

**Table S1:** Average component spectra of cells from glass substrates (n=14) and after Latrunculin-A (n=18). The Amide I band was decomposed by fitting Lorentzians (as explained in the main text) with known center wavenumber for each secondary structure.

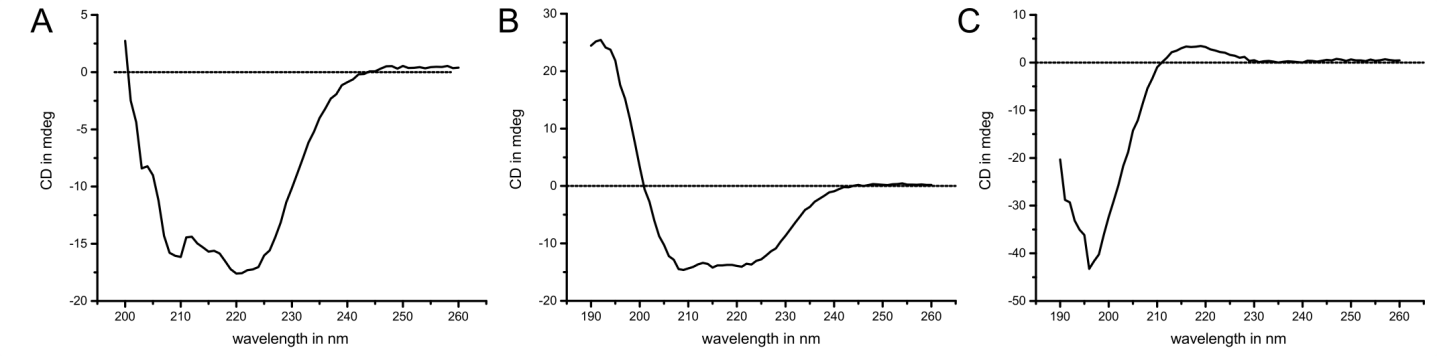

**Figure S17:** Circular dichroism spectroscopy of vimentin monomers in PBS showing the dominantly a-helical structure of the protein.

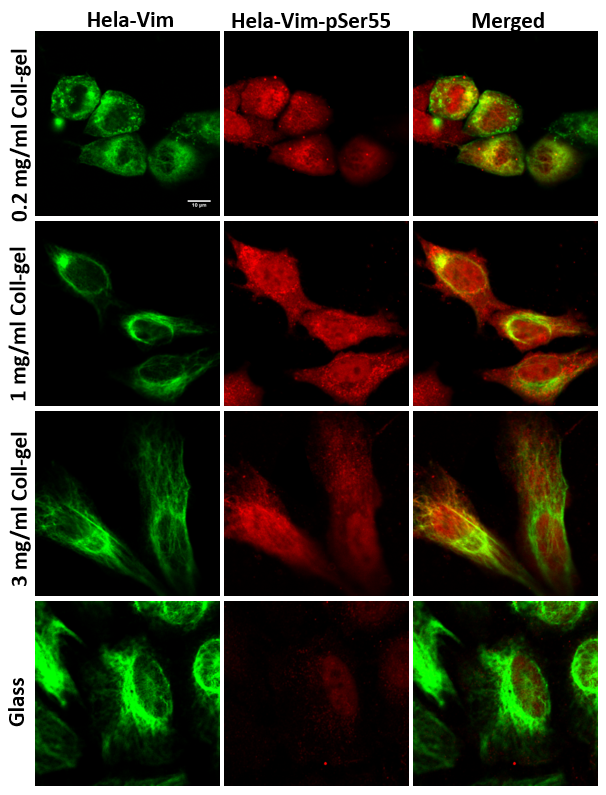

**Figure S18**: Immunofluorescence imaging of HeLa-GFPvim cells grown on soft collagen hydrogels (0.2 mg/ml, 1 mg/ml and 3 mg/ml) and glass surfaces shows GFP-tagged vimentin, pSer55-antibody staining and merged images. Scale bars are 10 µm.

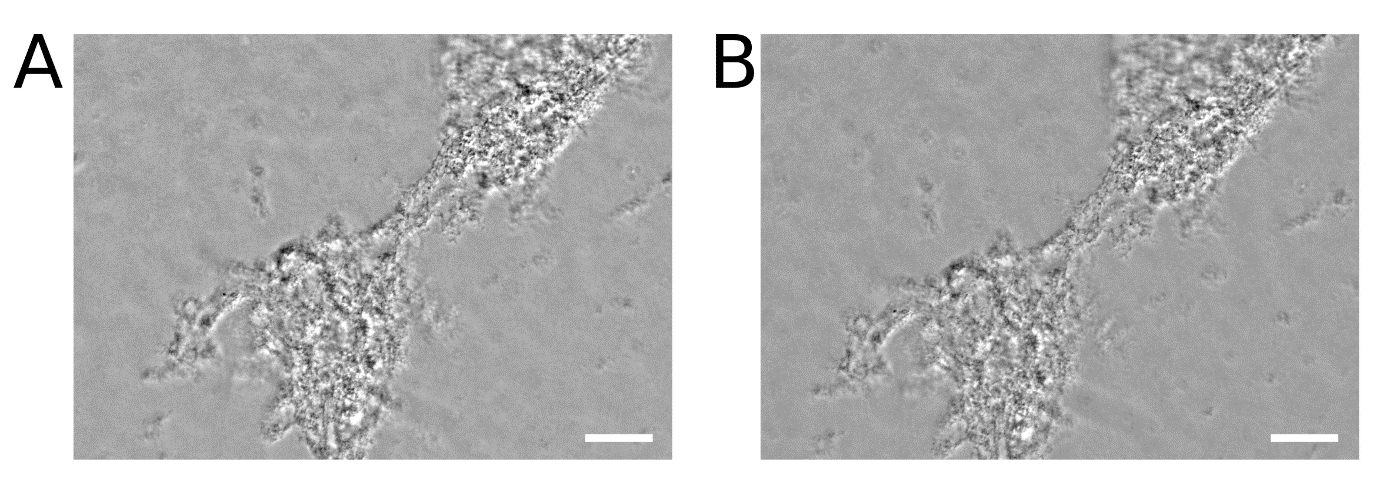

**Figure S19**: Phase contrast microscopy images of vimentin filaments formed *in vitro* before (A) and after 12 h blebbistatin treatment (B) show very similar appearance. Scale bars indicate 50 µm and a gradient background was subtracted from both images.

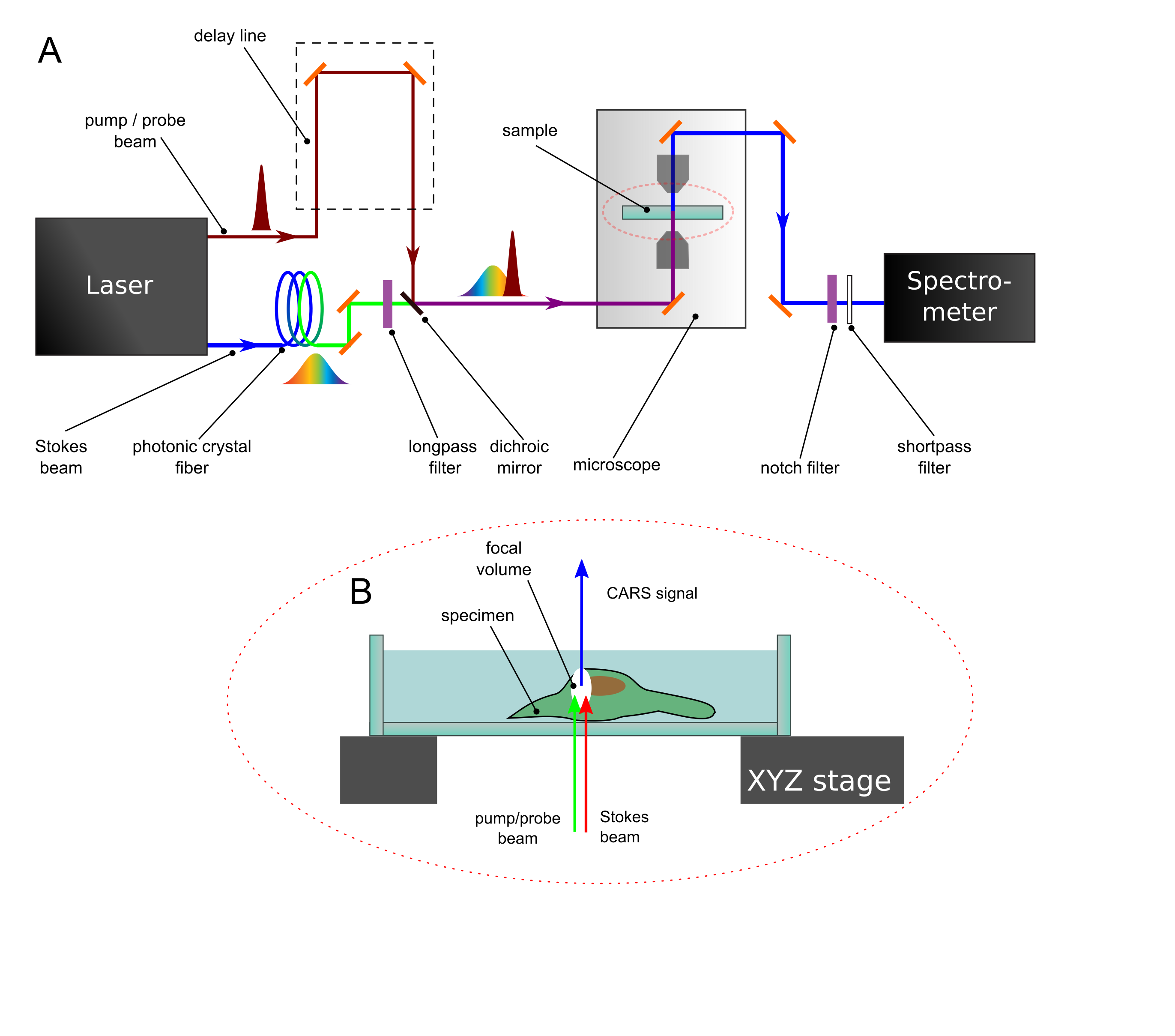

**Figure S20**: Illustration of the broadband CARS setup used in this work. A) Optical beam path for creating and detecting BCARS signal. B) Zoom-in showing the sample area where the Stokes beam is spatially and temporally overlapped with the pump/probe beam to create the forward-detected CARS signal.

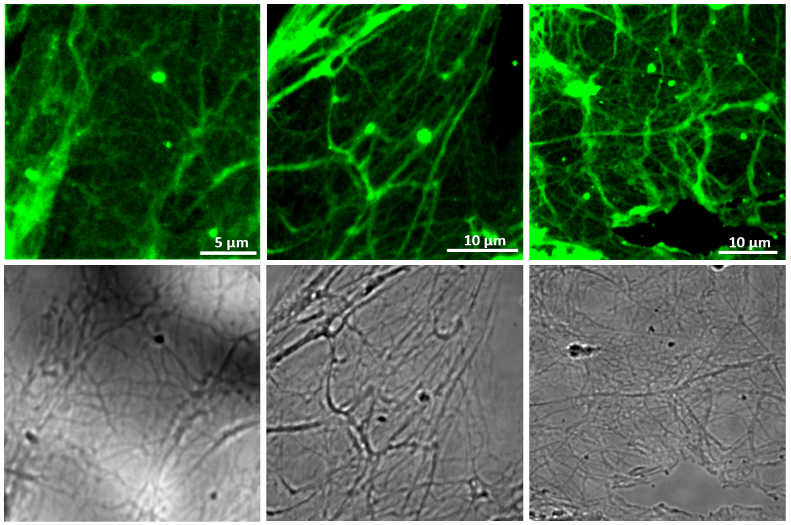

**Figure S21**: *In vitro* assembly of vimentin displaying networks of long and smooth filaments.
